# “The *B. subtilis* replicative polymerases bind the sliding clamp with different strengths to tune replication processivity and fidelity”

**DOI:** 10.1101/2025.03.10.642433

**Authors:** Luke G. O’Neal, Madeline N. Drucker, Ngoc Khanh Lai, Ashley F. Clemente, Alyssa P. Campbell, Lindsey E. Way, Sinwoo Hong, Emily E. Holmes, Sarah J. Rancic, Nicholas Sawyer, Xindan Wang, Elizabeth S. Thrall

**Affiliations:** Department of Chemistry and Biochemistry, Fordham University, Bronx, NY 10458; Department of Biology, Indiana University, Bloomington, IN 47405

**Keywords:** DNA replication, DNA polymerase, sliding clamp, single-molecule imaging, super-resolution microscopy

## Abstract

Ring-shaped sliding clamp proteins are essential components of the replication machinery, the replisome, across all domains of life. In bacteria, DNA polymerases bind the sliding clamp, DnaN, through conserved short peptide sequences called clamp-binding motifs. Clamp binding increases the processivity and rate of DNA synthesis and is generally required for polymerase activity. The current understanding of clamp-polymerase interactions was elucidated in the model bacterium *Escherichia coli*, which has a single replicative polymerase, Pol III. However, many bacteria have two essential replicative polymerases, such as PolC and DnaE in *Bacillus subtilis*. PolC performs the bulk of DNA synthesis whereas the error-prone DnaE only synthesizes short stretches of DNA on the lagging strand. How the clamp interacts with the two polymerases and coordinates their activity is unknown. We investigated this question by combining in vivo single-molecule fluorescence microscopy with biochemical and microbiological assays. We found that PolC-DnaN binding is essential for replication, although weakening the interaction is tolerated with only minimal effects. In contrast, the DnaE-DnaN interaction is dispensable for replication. Altering the clamp-binding strength of DnaE produces only subtle effects on DnaE cellular localization and dynamics, but it has a substantial impact on mutagenesis. Our results support a model in which DnaE acts distributively during replication but can be stabilized on the DNA template by clamp binding. This study provides new insights into the coordination of multiple replicative polymerases in bacteria and the role of the clamp in polymerase processivity, fidelity, and exchange.

## Introduction

Cellular DNA is replicated by a multi-protein complex called the replisome, the basic features of which are conserved from bacteria to humans (Figure 1A).(1–3) A central component of the replisome is the replicative DNA helicase, which unwinds the parental double-stranded DNA to generate single-stranded DNA (ssDNA). Another key component, the replicative DNA polymerase, catalyzes the synthesis of new DNA using the parental strands as templates. To synthesize the long stretches of DNA required to duplicate the genome, polymerases interact with sliding clamps, ring-shaped proteins that encircle the parental DNA and tether polymerases to the template.(4–6) DNA replication occurs by different mechanisms on the two parental DNA strands. On one, called the leading strand, the template DNA is oriented to allow continuous 5ʹ to 3ʹ synthesis of a new DNA strand. On the other, the lagging strand, synthesis occurs in short stretches of approximately 1 – 2 kilobases (kb) in length, known as Okazaki fragments.(1–3) Transiently exposed ssDNA on the lagging strand is bound by another replisome component, single-stranded DNA-binding protein (SSB), which plays several roles in replication, including protecting ssDNA from degradation.(7, 8) Finally, because replicative DNA polymerases cannot synthesize DNA de novo, each Okazaki fragment is initiated by synthesis of a short RNA primer by an enzyme known as the primase.(1–3)

**Figure 1.**
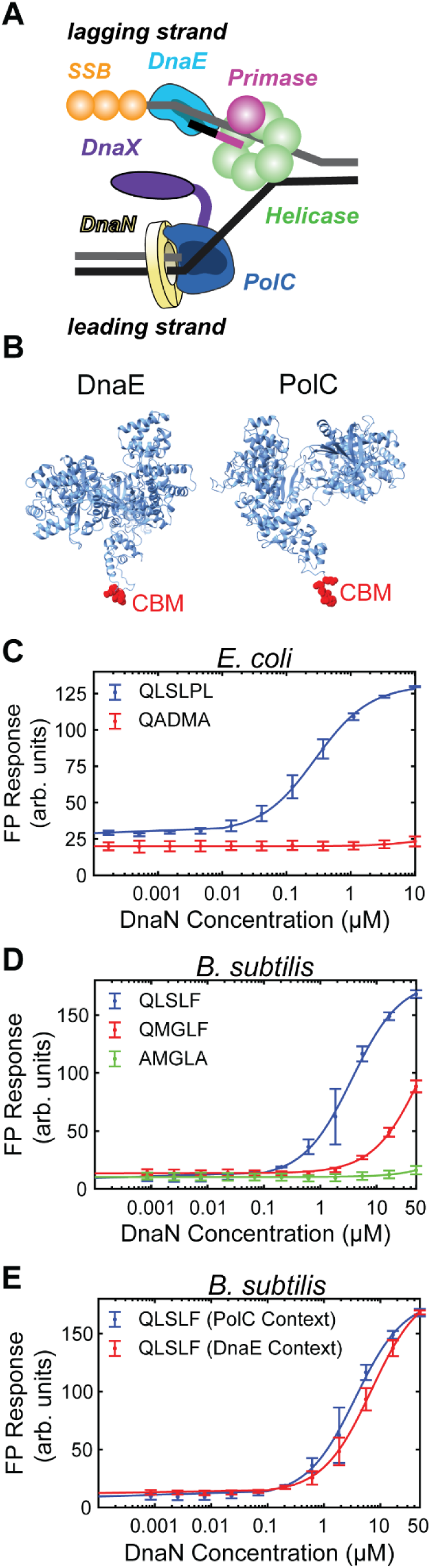
PolC and DnaE bind DnaN with different affinity. (A) Cartoon of the *B. subtilis* replisome. (B) AlphaFold predicted structures of *B. subtilis* DnaE (AlphaFold DB: AF-O34623- F1) and PolC (AlphaFold DB: AF-P13267-F1) with CBMs highlighted in red. See note on PolC structure in Figure S1. (B) FP response for *E. coli* DnaN with tight-binding (QLSLPL) and binding-deficient (QADMA) CBM peptides. (C) FP response for *B. subtilis* DnaN with PolC (QLSLF), DnaE (QMGLF), and binding-deficient (AMGLA) CBM peptides. (D) FP response for *B. subtilis* DnaN with PolC (QLSLF) CBM peptides with flanking amino acids from PolC or DnaE. Fits to a 1:1 binding model are represented as solid lines. Error bars show standard deviation of at least three replicates.

The molecular mechanisms of bacterial DNA replication are best understood in the gram-negative species *Escherichia coli*. In *E. coli*, replication is performed by a single replicative polymerase, Pol III, which contains polymerase (α) and exonuclease (ε) subunits as well an accessory subunit (θ).(1, 3) However, although the general replisome architecture is conserved across bacteria, it has become clear that there are key mechanistic differences in many species. In particular, many bacteria use more than one replicative polymerase.(3, 9) In low-GC gram-positive bacteria like the model organism *Bacillus subtilis*, both polymerases PolC and DnaE are essential for DNA replication and cell viability.(10–12) It was initially proposed that PolC and DnaE were the leading- and lagging-strand polymerases, respectively, by analogy to the eukaryotic polymerases Pol ε and Pol δ.(10) However, the rate of synthesis for DnaE is too slow to support the rate of replication fork progression observed in vivo.(11) In the current model, DnaE extends the RNA primer at the start of each Okazaki fragment, only synthesizing a short stretch of DNA before yielding to PolC, which performs the bulk of the synthesis on both strands.(11) This model implies frequent exchange between DnaE and PolC, and it is unclear how the *B. subtilis* replisome coordinates the activity of the two polymerases.

Compared to most replicative DNA polymerases, DnaE has several unusual features. Unlike PolC, it does not contain an integral exonuclease domain, nor has it been found to interact with an accessory exonuclease,(12, 13) analogous to the interaction between the α and ε subunits of *E. coli* Pol III. Thus, it is likely more error-prone than PolC. In the related low-GC gram-positive species *Streptococcus pyogenes*, the DnaE error rate was found to be two to three orders of magnitude higher than for *E. coli* Pol IIIα,(14) and depletion of DnaE in *B. subtilis* was found to reduce ultraviolet (UV)-induced mutagenesis.(13) Further, DnaE has the ability to bypass certain DNA lesions(13, 14) and is modestly upregulated via the SOS DNA damage response, a transcriptional program that increases the expression of DNA repair and damage tolerance factors, suggesting a possible role in DNA repair or damage tolerance.(13, 15)

To synthesize long stretches of DNA at high rates of ∼ 1,000 base pairs (bp) per second, replicative DNA polymerases must interact with processivity factors known as sliding clamps. In bacteria, the clamp is a homodimer that contains two identical binding sites, one on each protomer, for polymerases and other interacting proteins.(4–6) A variety of DNA replication and repair proteins bind to the sliding clamp at these sites through short amino acids segments called clamp-binding motifs (CBMs). The most common CBM is a pentapeptide with the consensus sequence QL[S/D]LF or more generally QxxLF;(12, 16) studies in *E. coli* have found that mutations to the conserved residues (particularly at the first, fourth, and fifth positions) significantly weaken or abolish clamp-binding.(16–18) Typical dissociation constants (*K*_D_) for the clamp-binding interaction in *E. coli* range from 100 nM – 1 μM.

Like all DNA polymerases in *E. coli*, both replicative polymerases in *B. subtilis* have canonical CBMs situated to allow clamp binding (Figures 1B and S1). The PolC CBM (QLSLF) is located at the protein C-terminus and is more conserved than the internal DnaE CBM (QMGLF). Given the presence of conserved CBMs, it is reasonable to assume that both PolC and DnaE interact with the clamp during replication. However, polymerase-clamp interactions in *B. subtilis* have never been quantified and the binding affinities are unknown. There is indirect evidence for PolC-DnaN binding in the form of a yeast two-hybrid assay,(19) and biochemical studies have shown that PolC activity is stimulated by the interaction with the clamp.(20) For DnaE, however, the results are conflicting. Although the same yeast two-hybrid assay did not find evidence for a DnaE-DnaN interaction,(19) several biochemical studies have shown that the presence of the clamp affects either the processivity or the rate of synthesis of DnaE.(11, 20, 21) Instead, other studies have proposed a role for DnaE interactions with other replisome components, including SSB,(21, 22) the clamp-loader complex subunit δ (HolA),(23) or a complex formed by the helicase DnaC and primase DnaG.(23)

In this study, we investigate the interaction between DnaN and the replicative polymerases PolC and DnaE in *B. subtilis*. We find that deleting the PolC CBM or mutating it to eliminate clamp-binding is lethal, but weakening the PolC-DnaN interaction by an order of magnitude is tolerated, though it leads to defects in DNA replication and changes in PolC cellular localization. Further, we show that DnaE also binds the clamp, albeit less tightly than PolC. Surprisingly, however, the DnaE-DnaN interaction is dispensable for DNA replication. Eliminating clamp-binding in DnaE has no effect on DNA replication, damage tolerance, or mutagenesis, nor does it alter the cellular localization and dynamics DnaE. Strengthening the DnaE-DnaN interaction also has no effect on DNA replication or damage tolerance, but it leads to a substantial increase in damage-induced mutagenesis and subtle changes in DnaE cellular localization and dynamics. These results provide new insights into the architecture of the *B. subtilis* replisome and the role of the sliding clamp in polymerase coordination in bacteria.

## Results

### Both the PolC and DnaE CBMs bind the clamp, but the DnaE CBM binds more weakly

Although PolC and DnaE contain canonical CBMs, no studies have quantitatively characterized their interaction with the clamp, DnaN. Previous studies in *E. coli* have shown that peptide CBMs are able to recapitulate clamp-binding using fluorescence polarization (FP),(24, 25) surface plasmon resonance (SPR),(17, 26) or isothermal titration calorimetry (ITC).(25, 26) Thus, we implemented an FP binding assay using purified DnaN and short peptides containing CBM sequences labeled with fluorescein (Figure S2 and Table S1). Similar to prior reports, we designed each peptide with 2 – 3 flanking amino acid residues from the native polymerase sequence on either end of the CBM as well as a 6-aminohexanoic acid residue linker and fluorescein appended to the N-terminus.

We first showed that we could reproduce prior clamp-binding measurements using purified *E. coli* DnaN (Figure 1C). We tested the binding of two peptides, one with a tight-binding CBM from the Hda protein (QLSLPL) and one with a binding-deficient CBM (QADMA), in which the terminal phenylalanine of the Pol IIIα CBM (QADMF) is mutated to alanine.(17) In a direct binding format, increasing concentrations of DnaN are incubated with a fixed concentration of fluorescently-labeled peptide and the FP response is measured. As expected, the tight-binding CBM binds to the clamp, showing a characteristic sigmoidal profile (Figure 1C). We fit the FP response vs. DnaN concentration to a 1:1 binding model to determine the dissociation constant, *K*_D_; although the dimeric clamp contains two equivalent binding sites, previous studies have used a 1:1 Langmuir model.(17, 26) Consistent with prior reports,(17, 27) we found that the tight-binding peptide binds *E. coli* DnaN with a sub-micromolar dissociation constant (mean ± std. dev.: *K*_D_ = 280 ± 90 nM), validating our FP assay design. As expected, the binding-deficient peptide showed no measurable binding up to a DnaN concentration of 10 µM (Figure 1C).

Next, we tested the binding of the wild-type (WT) PolC and DnaE CBM peptides to the *B. subtilis* clamp. Although we found that both CBM peptides bound the clamp, they did so with much lower affinity (Figure 1D). For the PolC CBM peptide, we measured *K*_D_ = 3.8 ± 1.8 μM, approximately 5-fold weaker than binding of the same CBM sequence to the *E. coli* clamp.(17) Although the DnaE CBM peptide also bound the clamp, it did so even more weakly than PolC, despite matching the consensus QxxLF motif. We did not test DnaN concentrations higher than 50 μM, but a fit to the data provided an approximate value of *K*_D_ = 78 ± 39 μM, roughly an order of magnitude weaker than PolC CBM binding and two orders of magnitude weaker than the binding of strong CBM sequences in *E. coli*. To test whether the measured binding was specific, we created a mutant DnaE CBM peptide with the highly conserved first and fifth residues mutated to alanine (AMGLA). Consistent with studies in *E. coli*, which found that mutating these residues substantially weakened binding,(16, 17) this double alanine DnaE CBM mutant showed minimal binding up to a DnaN concentration of 50 μM. Taken together, our measurements indicate that both PolC and DnaE can bind the *B. subtilis* clamp via their CBMs, albeit with significantly different affinities.

### The PolC-DnaN interaction is essential, but the DnaE-DnaN interaction is not

To examine whether the clamp interaction was essential for DNA replication, we first tested whether the PolC CBM could be eliminated or mutated by attempting to modify the *polC* gene. Because the PolC CBM is the last five amino acids at the C-terminus of the protein, we tried to truncate it. Despite multiple attempts, we were unable to recover transformants containing the truncation mutation, suggesting that the presence of the PolC CBM is essential. Consistent with the lack of binding observed for the AMGLA CBM peptide in the FP assays, we were likewise unable to recover transformants in which the PolC CBM was altered to AMGLA. Surprisingly, however, a PolC mutant (*polC^QMGLF^*) bearing the WT DnaE CBM, QMGLF, was viable, despite the CBM sequence having at least an order of magnitude lower affinity for the clamp than the WT PolC CBM. These results indicate that the PolC-DnaN interaction is essential to replication, but a significantly weaker clamp-binding interaction than the WT interaction still permits polymerase function.

Next, we tested whether the DnaE-DnaN interaction was likewise essential for DNA replication. Because the DnaE CBM is internal, we did not attempt to delete it; instead, we introduced the double-alanine AMGLA mutation, which was deficient in clamp binding in FP assays. Surprisingly, we found that a strain bearing this mutation (*dnaE^AMGLA^*) was viable. Thus, the DnaE-DnaN interaction is dispensable for DNA replication, in contrast to the essentiality of the PolC-DnaN interaction. In light of these results, we asked whether strengthening the DnaE- DnaN interaction would prove lethal by allowing DnaE to outcompete PolC for clamp binding. To test this question, we created a strain in which the DnaE CBM was mutated to the WT PolC CBM sequence QLSLF (*dnaE^QLSLF^*); binding of this PolC CBM sequence with the flanking amino acids from DnaE was similar to its binding in the native PolC context (*K*_D_ = 7.2 ± 2.2 μM) (Figure 1E). We found that a strain bearing the *dnaE^QLSLF^* mutation was viable, indicating that PolC could still gain access to the clamp in the presence of the tight-binding DnaE CBM mutant.

### Altering the DnaE-DnaN interaction does not affect cell growth or DNA replication, but altering the PolC-DnaN interaction does

After determining mutations in the PolC and DnaE CBMs that were tolerated in vivo, we next characterized the impact of these mutations on cell growth and DNA replication. First, we measured growth curves of the WT and CBM mutant strains in liquid culture. For growth in minimal S7_50_-sorbitol media, the WT strain has a doubling time of 60 ± 1 min (mean ± std. dev.) in exponential phase (Figure 2A and Table S2). We found no change in growth rate for the binding-deficient *dnaE^AMGLA^* mutant and the tight-binding *dnaE^QLSLF^* mutant (58 ± 1 min and 59.1 ± 0.6 min respectively; *p* > 0.05 vs. WT); the same was true for the *polC^QMGLF^* mutant (58 ± 2 min; *p* > 0.05). Reasoning that DNA replication defects might become more apparent in rich media, with faster cell growth, we also quantified the growth rate in LB Lennox media. Under these conditions, the WT doubling time is 31 ± 1 min (Figure 2B and Table S2). The *dnaE^QLSLF^* mutant was still indistinguishable from WT (30 ± 1 min; *p* > 0.05). Although the *dnaE^AMGLA^* doubling time was statistically distinguishable from WT, the change was minimal (28 ± 2 min; *p* < 0.05). However, the *polC^QMGLF^* mutant displayed a clear growth defect with a doubling time (41 ± 3 min; *p* < 0.05) that was 30% longer than WT.

**Figure 2.**
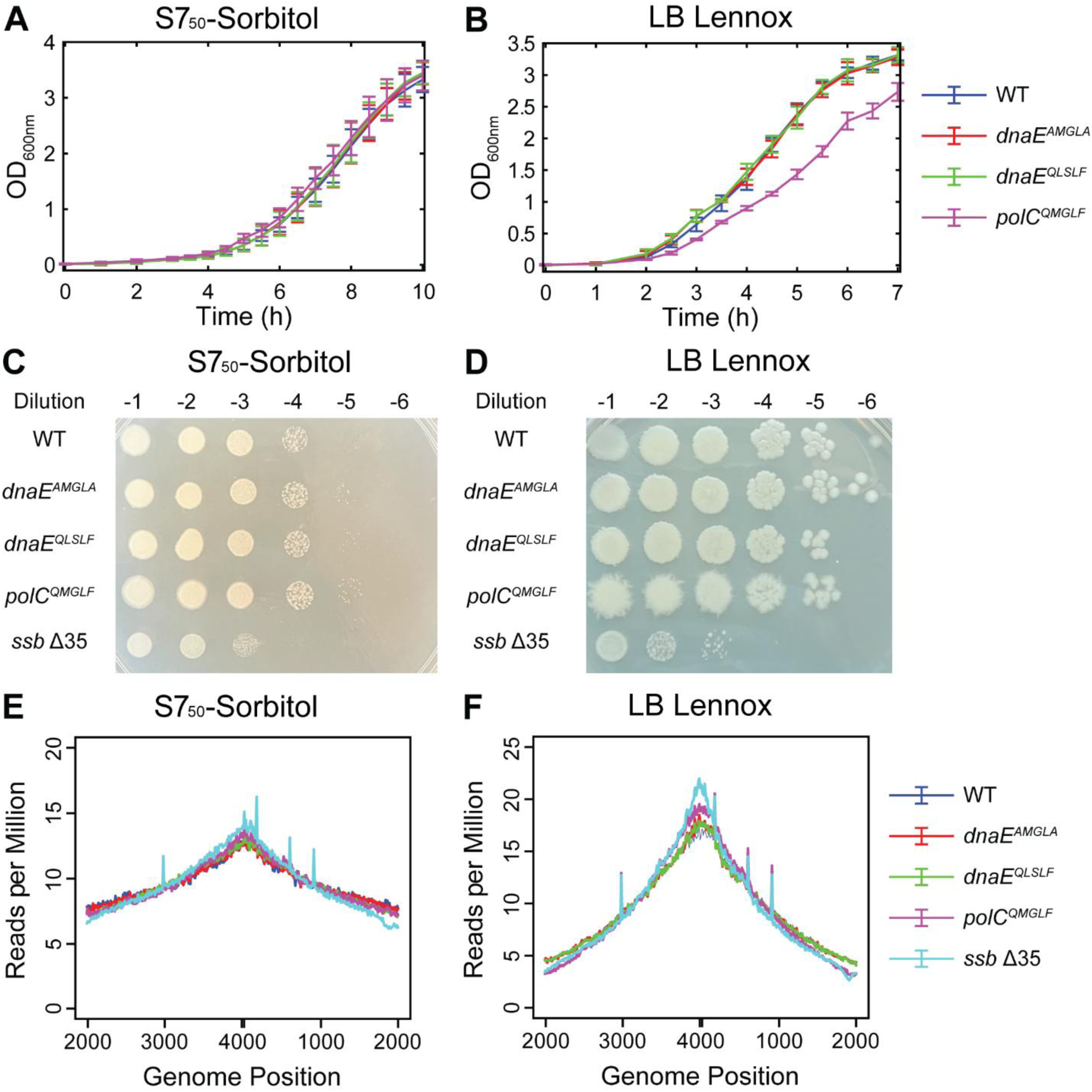
Growth and replication characterization of WT and CBM mutant DnaE and PolC strains. Liquid culture growth curves for WT, *dnaE^AMGLA^*, *dnaE^QLSLF^*, and *polC^QMGLF^* strains in (A) S7_50_- sorbitol and (B) LB Lennox media. Error bars show standard deviation of at least three replicates. (B) Serial dilution for WT, *dnaE^AMGLA^*, *dnaE^QLSLF^*, *polC^QMGLF^*, and *ssb* Δ35 strains on (C) S7_50_- sorbitol and (D) LB Lennox agar media. Whole-genome sequencing analysis showing the normalized number of reads vs. genome location for WT, *dnaE^AMGLA^*, *dnaE^QLSLF^*, *polC^QMGLF^*, and *ssb* Δ35 strains in (E) S7_50_-sorbitol and (F) LB Lennox media.

Because cell growth differs in liquid and solid media, we also characterized growth of the WT and CBM mutant strains on solid media. Cells were harvested from liquid cultures, serially diluted, spotted on agar plates containing media, and incubated overnight. The plates were analyzed qualitatively for differences in colony number and size. Consistent with the liquid culture growth curve results, growth of the DnaE and PolC CBM mutant strains was indistinguishable from that of the WT strain on minimal S7_50_-sorbitol agar (Figure 2C). Likewise, on rich LB Lennox agar, the DnaE CBM mutant strain growth was identical to WT (Figure 2D). Although the PolC CBM mutant strain did not display a viability defect, it displayed a spiky colony morphology. Because we did not observe a viability defect for any of the CBM mutant strains, we also tested a strain that was known to be significantly impaired in growth. This strain (*ssb* Δ35) bears a 35 amino acid truncation to the C-terminal of the replisome component SSB. Deletion of the SSB C-terminal tail was reported to confer substantial DNA replication and cell morphology defects.(22) Consistent with this prior report, we found that viability of the *ssb* Δ35 strain was reduced by roughly one or two orders of magnitude relative to WT in S7_50_-sorbitol and LB Lennox media, respectively, confirming that a substantial viability defect would be evident in these assays. Taken together with the liquid culture growth curves, these results indicate that DNA replication is moderately impaired when the PolC-DnaN interaction strength is reduced, although this effect is only obvious for fast growth in rich media. Surprisingly, however, we found no evidence that eliminating the DnaE-DnaN interaction alters cell growth.

To determine whether altering the DnaE-DnaN interaction led to DNA replication defects not apparent in growth assays, we analyzed genome-wide DNA replication profiles using marker frequency analysis (MFA) of whole-genome sequencing (WGS) results. Cultures were grown in S7_50_-sorbitol or LB Lennox media to early exponential phase, then cells were harvested and lysed. WGS (WGS) was performed, and the number of sequencing reads was plotted as a function of chromosomal position. Because replication initiates at the chromosomal origin (*ori*) and concludes at the terminus (*ter*), the DNA content and therefore relative number of reads is highest at *ori* and decays toward *ter*. Defects in DNA replication lead to changes in the DNA content profile and the ratio of *ori*:*ter* reads can be used as a single quantitative metric.(28, 29) As expected, the *ssb* Δ35 mutant had an altered replication profile and elevated *ori:ter* ratio relative to WT in both S7_50_- sorbitol and LB Lennox media (Figures 2E – F, Figure S3, and Table S3), consistent with a slowdown of replication.(29) However, we found that the binding-deficient *dnaE^AMGLA^* mutant and the tight-binding *dnaE^QLSLF^* mutant had essentially the same replication profiles as the WT strain in both growth media; the *polC^QMGLF^* mutant only showed altered replication profile under fast-growth conditions (LB Lennox media), with an elevated *ori:ter* ratio consistent with an elongation defect. Thus, altering the DnaE-DnaN interaction does not affect DNA replication, but altering the PolC-DnaN interaction slows down DNA replication.

### Strengthening the DnaE-DnaN interaction promotes damage-induced mutagenesis

Unusually for a replicative polymerase, DnaE is a member of the SOS regulon and is modestly upregulated upon DNA damage.(13, 15) It lacks an integral proofreading domain(12) and no accessory proofreading subunit, analogous to *E. coli* Pol IIIε, has been identified. Thus, it is more error-prone than *E. coli* Pol III and has also been shown to bypass certain DNA lesions more efficiently.(13, 14, 20) We therefore asked whether the clamp interaction might play a role in the response of DnaE to DNA damage.

First, we determined whether the DnaE-DnaN interaction had an effect on cell survival upon DNA damage. We tested two different DNA damaging agents, 4-nitroquinoline 1-oxide (4- NQO) and 254 nm UV light. 4-NQO generates various adenine and guanine lesions, particularly quinoline *N*^2^-dG adducts,(30–32) whereas short-wavelength UV-C light produces several highly blocking lesions, including cyclobutane pyrimidine dimers and 6-4 photoproducts.(33) Consistent with prior reports,(34, 35) we found that a strain lacking the translesion synthesis (TLS) polymerase Pol Y1 (Δ*yqjH*) was sensitized to both 4-NQO and UV treatment relative to the WT strain (Figures 3A – B). In contrast, both the binding-deficient *dnaE^AMGLA^* mutant and the tight-binding *dnaE^QLSLF^* mutant had WT levels of resistance to both types of DNA damage. Although these results do not exclude the possibility that an effect would be observed for other types of DNA lesions, they suggest that any involvement of DnaE in the DNA damage response does not require an interaction with the clamp, nor is it enhanced by a stronger clamp-binding interaction.

**Figure 3.**
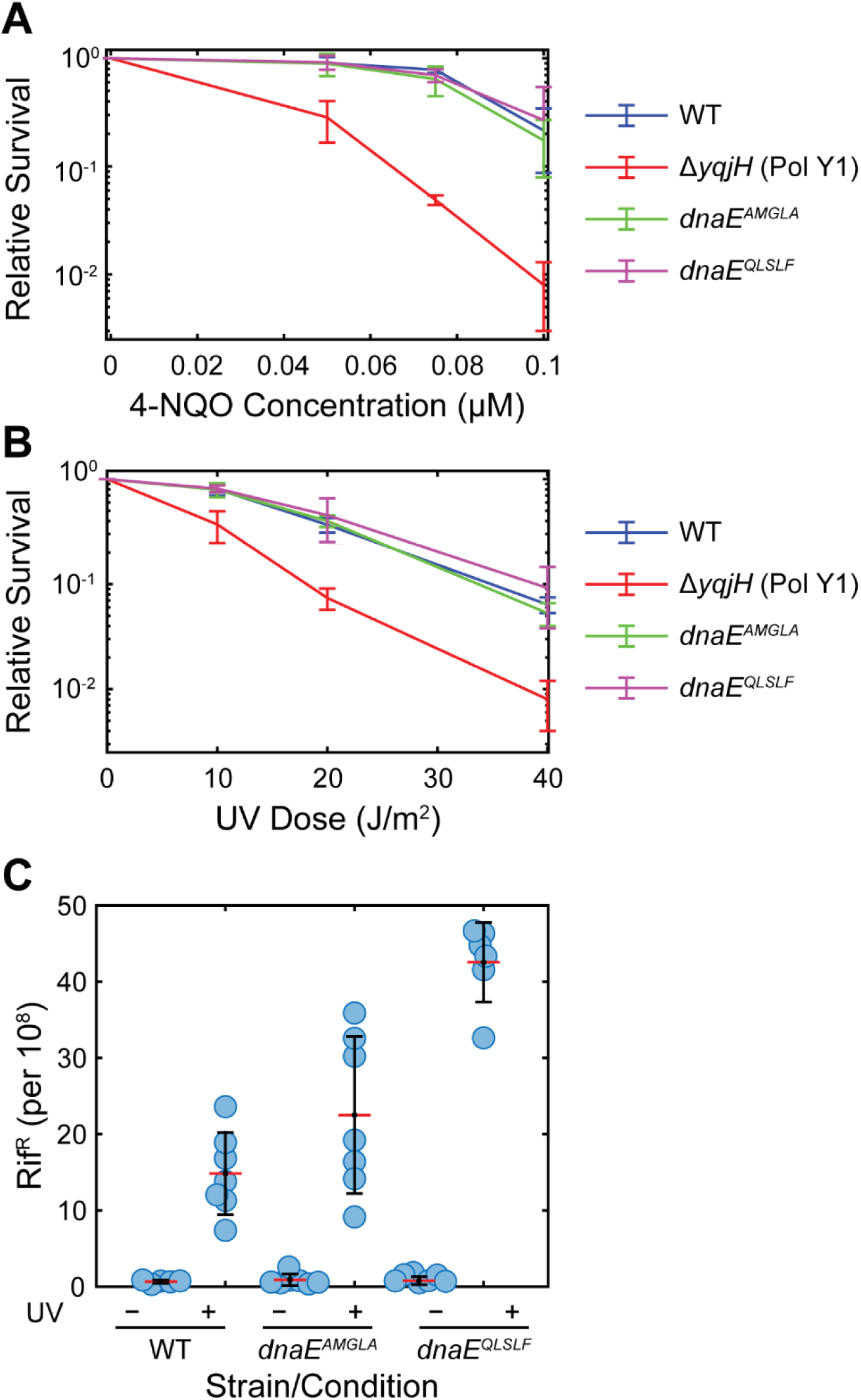
Relative survival and mutagenesis of WT and CBM mutant DnaE strains. (A) Relative survival of WT, *dnaE^AMGLA^ dnaE^QLSLF^*, and Pol Y1 knockout strains treated with different concentrations of 4-NQO. (B) Relative survival of WT, *dnaE^AMGLA^*, *dnaE^QLSLF^*, and Pol Y1 knockout strains treated with different doses of 254 nm UV light. For (A) and (B), error bars show standard deviation of at least three replicates. (C) Proportion of rifampicin resistant (Rif^R^) cells for WT, *dnaE^AMGLA^*, and *dnaE^QLSLF^* strains in untreated cells and cells treated with 40 J/m^2^ 254 nm UV light. For each dataset, the individual replicates are shown as circles, the red line represents the mean value, and the error bars represent the standard deviation.

Given the prior finding that DnaE contributes to UV-induced mutagenesis,(13) we next asked whether the DnaN interaction plays a role in mutagenesis. We measured the rate of resistance to a 10 μg/mL concentration of the drug rifampicin (Rif^R^) both in untreated cells and in cells treated with a 40 J/m^2^ dose of 254 nm UV light. This concentration of rifampicin is generally lethal, but mutations in the β-subunit of RNA polymerase (*rpoB*) can confer resistance,(36, 37) providing a proxy for the overall genomic mutation rate. Consistent with prior reports,(13, 35, 38) the spontaneous mutation rate for the WT strain was approximately one per 10^8^ cells (Figure 3C and Table S4). UV treatment increased this rate by approximately 20-fold (*p* < 0.05), again consistent with previous results. Both the *dnaE^AMGLA^* and *dnaE^QLSLF^* mutants had similar spontaneous mutation rates to WT (*p* > 0.05). However, the tight-binding *dnaE^QLSLF^* mutant had a 3-fold higher UV-induced mutation rate than the WT strain (*p* < 0.05). Although *dnaE^AMGLA^* showed a more modest increase in mutation rate relative to WT, the difference was not statistically significant (*p* > 0.05). These results support a model in which strengthening the DnaE-DnaN interaction allows DnaE to gain increased access to the DNA template upon perturbations to replication, increasing the mutation rate due to its error-prone activity.

### Altering the DnaN interaction has subtle effects on DnaE and PolC localization and dynamics in vivo

Taken together, our results support a model in which the PolC-DnaN interaction is essential for DNA replication; a binding affinity reduction of roughly an order of magnitude is tolerated, albeit with a growth defect in rich media. Although the DnaE-DnaN interaction is dispensable for replication and has no apparent effect on growth under any conditions tested, strengthening the interaction promotes mutagenesis, presumably by enhancing the access of DnaE to the DNA template. Next, we asked whether altering the clamp interaction had an effect on PolC and DnaE localization and dynamics in the cell using in vivo single-molecule fluorescence microscopy.

We created and validated an N-terminal fusion of the endogenous copy of DnaE to the self-labeling HaloTag (Halo);(39) we introduced an N-terminal tag because we had found previously that C-terminal fusions to the yellow fluorescent protein (YFP) variant mYPet(40) were impaired in growth, whereas N-terminal fusions were not (Figure S4B and Table S5). In addition to the Halo-DnaE fusion, we introduced a C-terminal mYPet fusion to the endogenous copy of the clamp-loader complex subunit DnaX to serve as a marker for replication sites in the cell (Figures 4A – B).(35, 41, 42) Growth curve measurements indicate that the Halo-DnaE fusion in combination with the DnaX-mYPet fusion has no effect on growth rate in the S7_50_-sorbitol media used for imaging (Figure S4A and Table S5), indicating that the fusions are functional. In addition, we were able to construct the corresponding binding-deficient Halo-DnaE^AMGLA^ and tight-binding Halo-DnaE^QLSLF^ fusions and to combine them with DnaX-mYPet.

**Figure 4.**
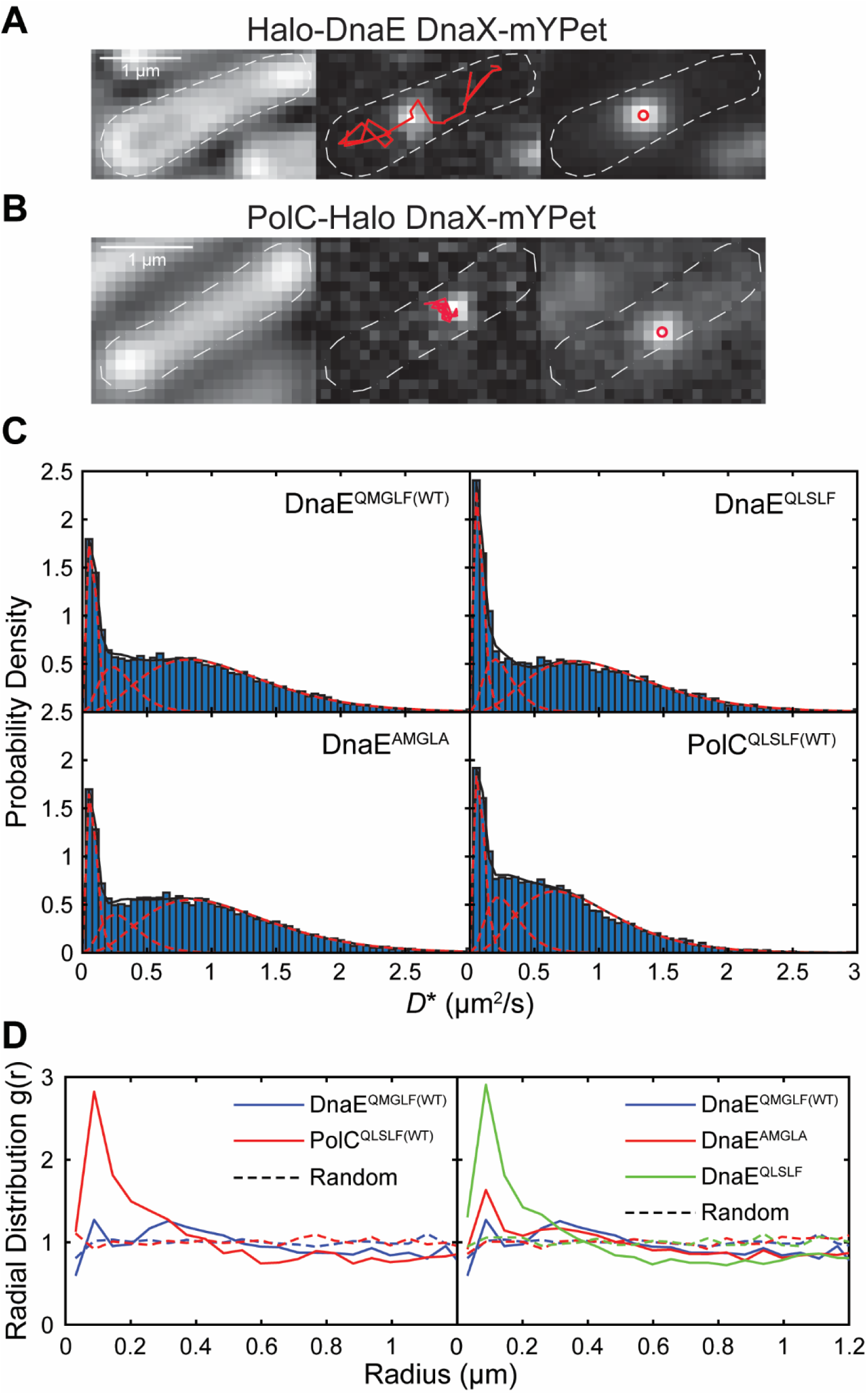
Imaging of single Halo-DnaE and PolC-Halo molecules. Representative micrographs of (A) Halo-DnaE^QMGLF(WT)^ with endogenous DnaX-mYPet fusion and (B) PolC^QLSLF(WT)^-Halo with ectopic DnaX-mYPet fusion. Left: transmitted white light micrograph of *B. subtilis* cell with overlaid cell outline and 1 μm scale bar. Middle: fluorescence micrograph of single Halo-DnaE^QMGLF(WT)^-JFX_554_ or PolC^QLSLF(WT)^-Halo-JFX_554_ molecule with overlaid trajectory. Right: fluorescence micrograph of DnaX-mYPet foci with overlaid centroids. (C) Apparent diffusion coefficient (*D*\*) distributions and corresponding three-population fits for Halo-DnaE^QMGLF(WT)^, Halo-DnaE^AMGLA^, Halo-DnaE^QLSLF^, and PolC^QLSLF(WT)^-Halo. (D) Radial distribution function *g*(*r*) analysis of Halo-DnaE or PolC-Halo colocalization with DnaX-mYPet. Left: Halo-DnaE^QMGLF(WT)^ and PolC^QLSLF(WT)^-Halo. Right: Halo-DnaE^QMGLF(WT)^, Halo-DnaE^AMGLA^, and Halo-DnaE^QLSLF^. Random *g*(*r*) curves are shown as dashed lines. *g*(*r*) values > 1 indicate colocalization relative to chance.

Likewise, we created a C-terminal HaloTag fusion to the endogenous copy of PolC. We found that this fusion was synthetically lethal with the endogenous DnaX-mYPet fusion, possibly due to an interaction between PolC and the DnaX C-terminus.(19, 20) However, we were able to introduce the DnaX-mYPet fusion at an ectopic locus (*yvbJ*) in combination with PolC-Halo; presumably clamp-loader complexes in this merodiploid strain contain a mix of tagged and untagged DnaX molecules, restoring viability. This strategy has been used with other PolC and DnaX fusions in previous studies.(43, 44) Growth curve measurements showed that this strain did not have a growth defect in S7_50_-sorbitol (Figure S4A and Table S5), confirming functionality of the fusions. Unfortunately, we were unable to make the corresponding weak-binding PolC^QMGLF^- Halo fusion; presumably the presence of the HaloTag at the PolC C-terminus in combination with the weaker CBM was too great of an impairment. We tried restoring functionality by using a longer 30 amino acid linker between the PolC C-terminus and the HaloTag, but we were still unable to recover the correct CBM mutant transformants.

Next, we validated a strategy for imaging single DnaE and PolC molecules in the cell. We used sparse labeling with low concentrations (1 nM and 500 pM, respectively) of the bright and photostable JFX_554_ dye(45) conjugated to a HaloTag ligand (Figures 4A – B). We confirmed that the labeling procedure did not affect cell growth by comparing the change in the number of colony-forming units (CFUs) per mL in labeled vs. mock-labeled cultures (Table S6).(35) We also verified that there was negligible detection of false positive spots in the JFX_554_ channel by labeling and imaging a strain bearing the DnaX-mYPet fusion alone (Figure S5).

After establishing the imaging approach, we first looked for changes in the DnaX foci or cell morphology due to DnaE CBM mutations. As a core component of the replisome, DnaX forms distinct foci at sites of replication.(35, 41, 42) In cells bearing the endogenous DnaX-mYPet fusion alone, there were typically one or two DnaX foci per cell (mean ± S.E.M.: 1.62 ± 0.03) with average localization at quarter and three-quarter positions along the long cell axis and at midcell along the short cell axis (Figure S6A), in good agreement with previous results.(35) There were minimal changes (5% or less) in the number of DnaX foci per cell in cells bearing the Halo-DnaE^QMGLF(WT)^ fusion, the binding-deficient Halo-DnaE^AMGLA^ fusion, and the tight-binding Halo-DnaE^QLSLF^ fusion (Table S7). Likewise, there were no qualitative changes in the cellular localization of the DnaX foci in these strains (Figure S6D – F). We characterized cell morphology by measuring the average length and width of cells and again found only minor differences in the presence of the WT Halo-DnaE fusion or the corresponding CBM mutant fusions (Table S7). Taken together, these results show no obvious perturbations to cell morphology or replisome positioning for the DnaE CBM mutants, consistent with our growth and replication assays.

Although the presence of the Halo-DnaE fusion appeared to have no substantial effects on DnaX or cell morphology, the PolC-Halo fusion was not as well tolerated. The cellular localization of the ectopic DnaX-mYPet fusion alone looked similar to that of the endogenous fusion (Figure S6B), although there was a 15% reduction in the number of foci per cell (mean ± S.E.M.: 1.39 ± 0.03; *p* < 0.05) (Table S7), possibly due to missed detection of dimmer foci that contain a mix of labeled and unlabeled DnaX molecules. In the presence of the PolC-Halo fusion, however, the DnaX localization shifted toward midcell along the long cell axis (Figure S6C); similar shifts in replisome positioning were observed previously in the presence of DNA damage,(35) suggesting a mild replication defect. In addition, there were subtle changes in cell morphology, with an increase in the average cell length of approximately 27% (*p* < 0.05) relative to the strain with the ectopic DnaX-mYPet fusion alone (Table S7). Thus, simultaneous labeling of PolC and DnaX appeared to confer mild growth defects, even though growth curve measurements did not reveal any changes.

Next, we used single-particle tracking to characterize the mobility of single WT DnaE and PolC molecules in the cell by calculating an apparent diffusion coefficient (*D*\*) from the mean squared displacement (MSD) of each trajectory.(35, 46) Consistent with prior studies of DNA polymerases in *B. subtilis*(35) and *E. coli*,(46, 47) we observed both static (*D*\* ≈ 0) and mobile populations of each polymerase (Figure 4C); the former population is expected to be bound to DNA or to DNA-bound proteins, although we cannot distinguish molecules that are synthesizing DNA from those that are not. To extract diffusion coefficients and the percentage of molecules in each population, we fit the *D*\* distributions to a three-population model because we found that a two-population model did not fit the data well. For both DnaE and PolC, there was a static population at *D*\* ≈ 0.08 μm^2^/s corresponding to 18% and 19% of the total population, respectively, as well as a population with intermediate mobility (*D*\* ≈ 0.30 μm^2^/s, 16% and 19% of the total population respectively) (Table S8). The largest population had higher mobility, with *D*\* ≈ 1.10 μm^2^/s (67%) for DnaE and *D*\* ≈ 0.88 μm^2^/s (62%) for PolC. Consistent with this analysis, and in agreement with an alternative analysis of mobility using the Spot-On package (Figure S7 and Table S9),(48) we found that the DnaE mobile population is broader and shifted to larger *D*\* values than the PolC mobile population. Consistent with these measurements, DnaE molecules visually appeared more mobile in the cell, as reported previously,(43) and their average localization in the cell was more diffuse (Figure S8).

Having characterized the mobility of DnaE^QMGLF(WT)^ molecules, we imaged the corresponding CBM mutant fusions to determine whether altering the clamp interaction had an effect on DnaE mobility. Both the binding-deficient DnaE^AMGLA^ and tight-binding DnaE^QLSLF^ mutants had similar overall *D*\* distributions to DnaE^QMGLF(WT)^, with similar *D*\* values for the static and higher mobility populations (Figure 4C and Table S8). However, there was a minor reduction in the static population for the DnaE^AMGLA^ mutant relative to DnaE^QMGLF(WT)^ (16% vs. 18%) and a likewise minor increase in the static population for the DnaE^QLSLF^ (21% vs. 18%); analysis results with the alternative Spot-On approach were in qualitative agreement (Table S9). These results are consistent with a model in which DnaE is not immobilized via interactions with the clamp during normal replication, and thus the change in static population is modest for the binding-deficient mutant. However, our data indicate that DnaE is capable of interacting with the clamp when the clamp-binding interaction is strengthened, leading to an increase in static population for the tight-binding mutant.

We also asked whether the clamp interaction plays a role in enriching DnaE at the replication fork by measuring polymerase colocalization with replication sites marked by DnaX-mYPet foci. To quantify colocalization, we used radial distribution function analysis, which provides more information on intracellular colocalization than does a comparison of average cellular localization. In brief, the radial distribution function *g*(*r*) gives a measure of the fold-enrichment of DnaE or PolC as a function of distance from the closest DnaX focus relative to the apparent enrichment that would be observed by chance due to confinement in the cell.(35, 47, 49) No enrichment relative to chance corresponds to *g*(*r*) = 1 and larger values of *g*(*r*) indicate greater enrichment. Consistent with the expectation that PolC performs processive synthesis at the replication fork whereas DnaE only synthesizes short stretches of DNA, we observed moderate PolC-DnaX colocalization (maximum *g*(*r*) ≈ 2.8) and little to no DnaE-DnaX colocalization (maximum *g*(*r*) ≈ 1.3) (Figure 4D and Table S10). The binding-deficient DnaE^AMGLA^ mutant had a similar degree of colocalization with DnaX (maximum *g*(*r*) ≈ 1.6) as DnaE^QMGLF(WT)^. However, the tight-binding DnaE^QLSLF^ mutant showed a clear increase in colocalization with DnaX (maximum *g*(*r*) ≈ 2.9), with a maximum value of *g*(*r*) similar to that of PolC^QLSLF(WT)^. These results are consistent with the DnaE diffusion measurements, suggesting that clamp binding has little to no role in stabilizing DnaE at the replication fork in WT cells, although strengthening the clamp interaction can promote DnaE binding at the replication fork thereby increasing its access to the DNA template.

Finally, we assessed the effect of weakening the PolC CBM at the cellular level. Although we were unable to create a viable HaloTag fusion to the weak-binding PolC^QMGLF^ mutant, we found that the corresponding PolC^QMGLF^-mYPet fusion was functional (Figure S4C and Table S5). Thus, we characterized the PolC-mYPet foci (Figures 5A – B) that form at replication sites to determine the effect of the CBM mutation. First, we measured the morphology of cells bearing the PolC^QLSLF(WT)^-mYPet and PolC^QMGLF^-mYPet fusions and found that there was a ∼ 22% increase (*p* < 0.05) in cell length for the CBM mutant (mean ± S.E.M.: 4.36 ± 0.04 μm) relative to WT (3.58 ± 0.04 μm) (Table S7). Next, we measured the average number of PolC foci per cell; we found no difference between PolC^QLSLF(WT)^ and the PolC^QMGLF^ mutant (mean ± S.E.M: 1.71 ± 0.04 vs. 1.78 ± 0.04; *p* > 0.05) (Table S7). However, we found changes in the average cellular localization of PolC foci for the WT and CBM mutant (Figure 5C). Consistent with the normal replication fork positioning observed for DnaX-mYPet foci, PolC^QLSLF(WT)^ formed foci at the quarter and three-quarter positions along the long cell axis. Conversely, the localization of PolC^QMGLF^ foci peaked at midcell along the long cell axis. As discussed previously, this change in replication fork positioning is indicative of perturbations to replication and is consistent with the defects observed in growth and replication assays for this weak-binding PolC CBM mutant.

**Figure 5.**
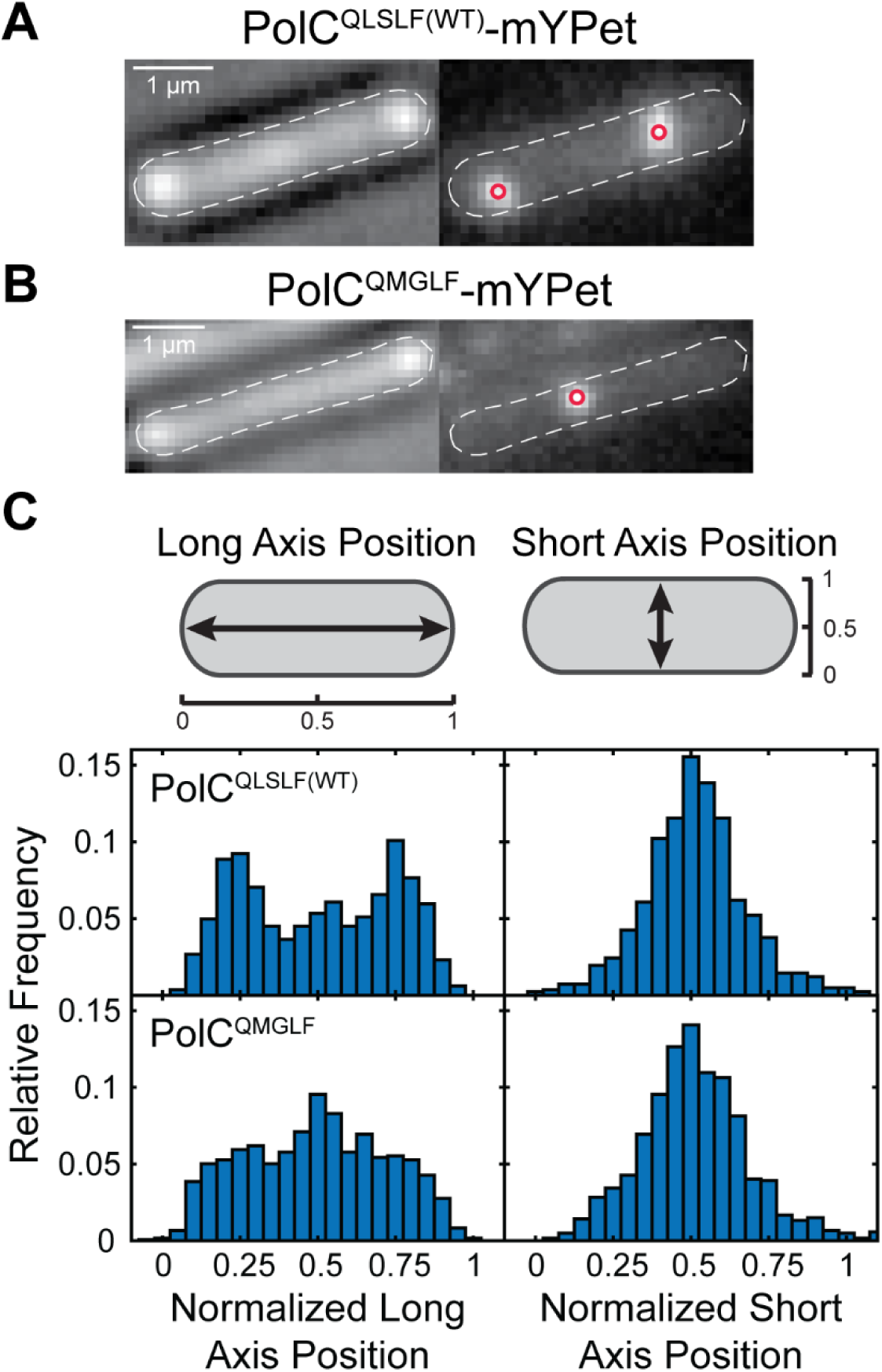
Imaging of PolC-mYPet foci. Representative micrographs of (A) PolC^QLSLF(WT)^-mYPet and (B) PolC^QMGLF^-mYPet foci. Left: transmitted white light micrograph of *B. subtilis* cell with overlaid cell outline and 1 μm scale bar. Right: fluorescence micrograph of PolC-mYPet foci with overlaid centroids. (C) Average cellular localization projected along normalized long and short cell axes for PolC^QLSLF(WT)^-mYPet and PolC^QMGLF^-mYPet foci.

## Discussion

In this study, we investigated the role of the sliding clamp interaction in coordinating replicative polymerase activity in *B. subtilis*. Both PolC and DnaE contain conserved CBM sequences, and we found that these CBM peptides are capable of binding the DnaN clamp. Although the PolC CBM binds the clamp with moderate affinity (mid-micromolar range), the DnaE CBM binds very weakly (*K*_D_ > 50 μM). By introducing CBM mutations to the chromosomal copies of PolC and DnaE and characterizing the impact on growth and replication, we found that the DnaE-DnaN interaction is dispensable for replication in *B. subtilis*; either strengthening or eliminating this interaction had no effect under any conditions tested. In contrast, the PolC-DnaN interaction is essential, although it can be weakened by roughly an order of magnitude with modest effects on growth and replication apparent in rich media only.

Results from quantitative in vivo fluorescence microscopy are consistent with this model. We found that altering the DnaE-DnaN interaction strength has no effect on cell morphology or on the localization of the replisome marked by DnaX, consistent with our characterization of growth and replication in DnaE CBM mutant strains. However, strengthening the DnaE-DnaN interaction leads to an increase in the static, or bound, population of DnaE molecules and a modest increase in DnaE colocalization with sites of replication. These results support the idea that DnaE is capable of binding the clamp when the CBM is strengthened, consistent with our finding that there is an increase in UV-induced mutagenesis for the tight-binding DnaE^QLSLF^ mutant. In contrast, weakening the PolC-DnaN interaction strength leads to measurable changes in cell morphology and the cellular localization of replication sites.

Our results support a model in which DnaE acts distributively during replication, consistent with biochemical evidence that DnaE only adds a small number of nucleotides to the RNA primer before PolC takes over synthesis (Figure 6).(11) However, it is possible that other protein-protein interactions play a role in coordinating DnaE activity during replication. DnaE interacts with SSB at a common binding site on the SSB C-terminal tail.(21, 22) Truncation of the SSB C-terminal tail, which is lethal in *E. coli*,(50) is tolerated in *B. subtilis*, albeit with defects in replication and cell morphology.(22) DnaE foci observed in WT cells were lost in SSB C-terminal tail truncation mutants, suggesting that this interaction may play a role in replication. Because the SSB binding site on DnaE is unknown and the SSB C-terminal tail truncation mutation is pleiotropic, it is unclear whether these defects arise partially or entirely from an impairment of DnaE activity or as a consequence of perturbing the interaction of other proteins with SSB. In addition, yeast two-hybrid and gel filtration chromatography showed that DnaE forms a ternary complex with DnaC helicase and DnaG primase, consistent with a model in which this complex recruits DnaE to the lagging strand and coordinates primer handoff from DnaG to DnaE.(23) Further investigation of these interactions will provide a more complete understanding of DnaE activity in replication in *B. subtilis*.

**Figure 6.**
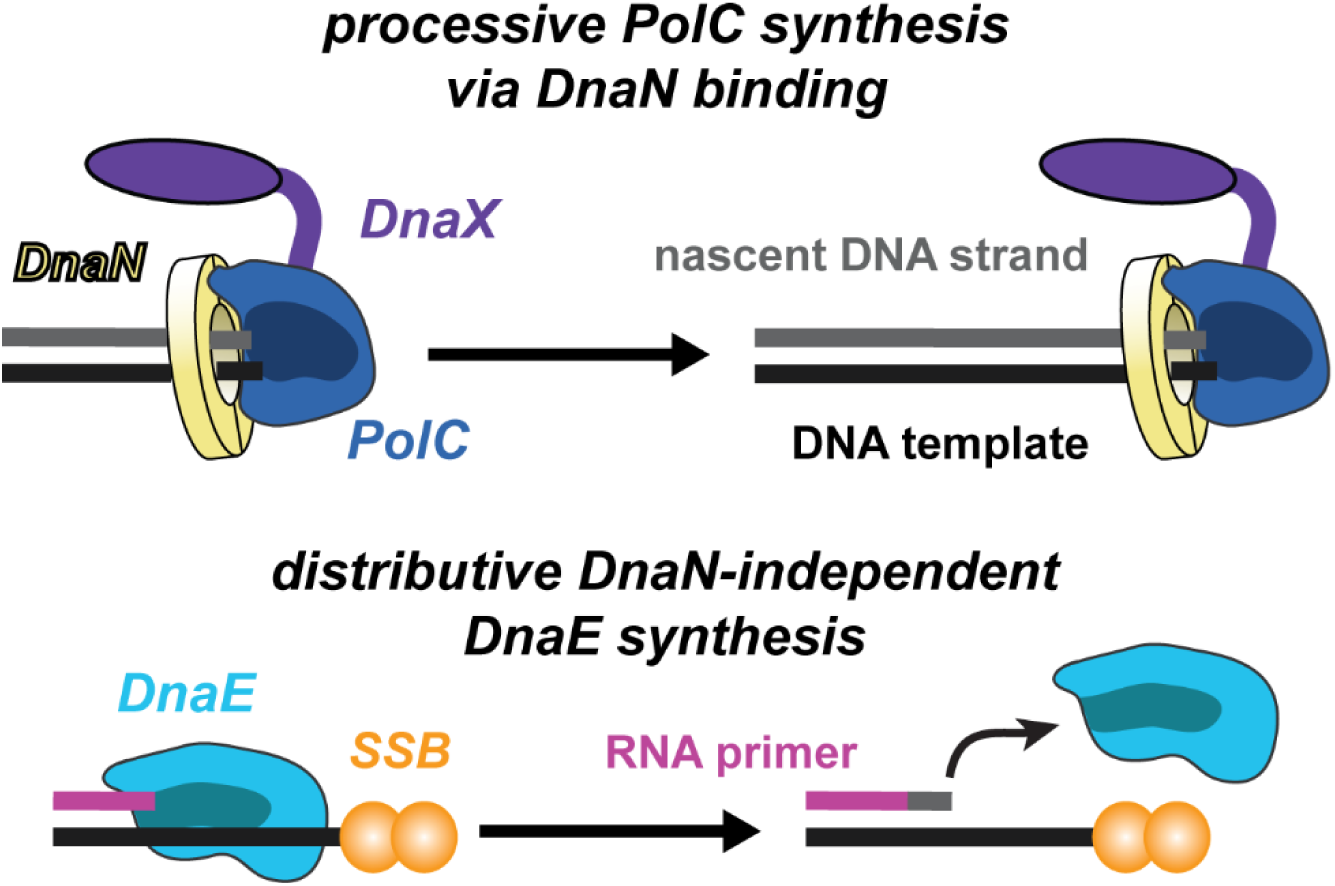
Cartoon of PolC and DnaE activity. Processive synthesis by PolC via the clamp interaction (top) contrasted with distributive clamp-independent synthesis by DnaE (bottom).

Although we have focused on *B. subtilis*, our results have implications for replication in other bacterial species. Phylogenetic analysis of different bacterial C-family polymerases has identified four broad categories, designated as PolC and three different DnaE subtypes (DnaE1, DnaE2, and DnaE3).(12) *E. coli* DnaE was assigned to the DnaE1 subtype and *B. subtilis* DnaE to the DnaE3 subtype. DnaE3 polymerases are never the sole replicative polymerase but are always present in combination with a PolC polymerase. The canonical CBM is almost always present and highly conserved across PolC proteins (QLSLF), and is likewise present across DnaE1 proteins, though with less conservation (QxxLF). However, only 75% of DnaE3 proteins contain an identifiable CBM, and there is substantial variability in the first position, with only the fourth and fifth positions conserved (xxxLF); thus, *B. subtilis* DnaE is unusual among DnaE3 proteins in having a more canonical CBM with a glutamine in the first position. Further, glycine is the most common residue in the third position across DnaE3 proteins,(12) which contrasts with the consensus aspartate or serine for all clamp-binding motifs. Glycine at the third position was shown to significantly weaken clamp binding in *E. coli*,(25) possibly due to an entropic binding penalty. Taken together with this phylogenetic analysis, our results suggest that DnaE polymerases across other bacterial species likewise do not require the clamp interaction for replication, and that the strength of the DnaE CBM may be tuned to avoid competition with PolC.

Nonetheless, our results support a model in which CBM strength alone is not the only determinant of clamp binding and polymerase selection. The viability of the weak-binding *polC^QMGLF^* mutant strain indicates that a range of CBM strengths can be tolerated by replicative polymerases, although it is unclear over what range. A prior study in *E. coli* measured the binding affinity of different Pol IIIα CBM mutants and assessed their ability to complement a temperature-sensitive Pol IIIα mutant when expressed at different levels in vivo.(51) Mutations that weakened the WT CBM (QADMF; *K*_D_ = 0.8 μM) by up to 5-fold (for example, AADMF; *K*_D_ = 4.2 μM) or strengthened it (QLDLF; *K*_D_ ≈ 6.9 nM) were tolerated, whereas 10-fold or higher reductions in affinity (for example, QADMK; *K*_D_ = 12.6 μM) were not. Conversely, the Pol IIIε subunit, which binds very weakly to the clamp (QTSMAF; *K*_D_ = 210 μM) could be weakened by more than 10- fold (ε_Q_: ATSMAF; *K*_D_ > 2 mM) or strengthened by almost 1000-fold (ε_L_: QLSLPL; *K*_D_ = 0.38 μM) with no effect on viability, although with some effects on the DNA damage response and mutagenesis.(27, 52, 53) In this study, we observed that a 10-fold reduction in binding affinity for the *polC^QMGLF^* mutant produced only modest phenotypes, suggesting that further reductions in affinity might be tolerated. An overly strong DnaE CBM could lead to replication defects if it allowed DnaE to outcompete PolC for access to the clamp. In *E. coli*, the interaction between Pol IIIα and the τ subunit of the clamp-loader complex helps to chaperone Pol III onto newly-loaded clamps,(54) but it is unknown whether a similar mechanism plays a role in polymerase selection in *B. subtilis*.

One surprising implication of our study is that clamp-binding interactions may be generally weaker in *B. subtilis* than in *E. coli*. Although extensive biochemical work has characterized the molecular determinants of clamp-binding in *E. coli*, much less is known in other bacteria. In general, the tightest CBM interactions are on the order of 200 – 400 nM for *E. coli* DnaN.(17, 25) Thus, it is surprising that a canonical CBM motif (QLSLF) binds *B. subtilis* DnaN with only micromolar affinity, whereas the same CBM was found to have submicromolar affinity in *E. coli*.(17) It is unknown whether this roughly order of magnitude difference in affinity holds for other CBM sequences in *B. subtilis* and, if so, whether there are functional reasons or consequences. Crystal structures have revealed differences in the *B. subtilis* DnaN binding site relative to *E. coli* that may contribute to weaker CBM binding in *B. subtilis*,(26, 55) although a structure of a peptide-bound complex with *B. subtilis* DnaN would be necessary to draw definitive conclusions.

Finally, our study provides new insights into the role of DnaE in the DNA damage response and damage-induced mutagenesis. The observation that DnaE is modestly SOS-inducible and mutagenic suggests that it may have a role in the DNA damage response, whether through a role in repriming or by acting as a translesion polymerase to bypass DNA lesions. However, we found that altering the DnaE-DnaN interaction had no effect on cell survival upon UV or 4-NQO treatment. In *E. coli*, one pathway to alleviate replication stalling is repriming downstream of the lesion, and there is evidence that the replisome alone is capable of repriming without other factors.(56) Thus, if DnaE plays a role in repriming after replication stalling in *B. subtilis*, which seems likely given the inability of PolC to extend an RNA primer,(11) this activity does not require an interaction with the clamp. Surprisingly, however, we found that strengthening the DnaE-DnaN interaction led to a substantial increase in UV-induced mutagenesis, but no change in spontaneous mutagenesis. These results are consistent with a model in which DnaE is able to gain access to DNA template via stronger clamp-binding interaction only when replication is perturbed. The lack of an effect on spontaneous mutagenesis suggests tight regulation of clamp binding at the replication fork, again consistent with the model that CBM strength is not the only determinant of polymerase selection. These results are also analogous to findings that the presence or absence of translesion polymerases only affects the damage-induced mutation rate and not the spontaneous mutation rate in both *E. coli*(57, 58) and *B. subtilis*.(38, 59) Further investigation is needed to understand the role that DnaE plays in the response to DNA damage how its mutagenic activity is limited during normal growth.

In summary, we found that the replicative polymerase DnaE acts independently of the clamp during DNA replication in *B. subtilis*. In contrast, clamp binding is essential for the activity of the replicative polymerase PolC. Strengthening the interaction between DnaE and the sliding clamp promotes mutagenesis, suggesting that the relative polymerase-clamp interaction strengths may be tuned to optimize replication processivity and fidelity. Our work has implications both for replisome organization in bacteria that use more than one replicative polymerase and for genome stability in species with error-prone replicative polymerases.

## Materials and methods

### Bacterial Strain and Plasmid Construction

Plasmids were constructed by amplifying double-stranded DNA (dsDNA) fragments using polymerase chain reaction (PCR), joining fragments by Gibson assembly,(60) and transformation into the cloning strain *E. coli* DH5α. Transformants were streak-purified once and then plasmids were miniprepped and validated by whole-plasmid sequencing. All *B. subtilis* strains were constructed in the wild-type (WT) PY79 background. All genetic modifications were introduced at the endogenous chromosomal loci unless noted otherwise. New genetic modifications were created by transformation of dsDNA fragments amplified by PCR and joined by Gibson assembly. Genetic modifications were transferred between strains by transformation of genomic DNA. Genetic modifications were linked to antibiotic resistance cassettes and transformants were selected on plates containing the corresponding antibiotic. Modified genetic loci were validated by diagnostic PCR and Sanger DNA sequencing. All oligonucleotides, plasmids, and bacterial strains used in this study are listed in Tables S11 – S13.

### Protein Expression and Purification

pET-28b plasmids expressing N-terminally hexahistidine (His_6_)-tagged *E. coli* and *B. subtilis* DnaN were transformed into *E. coli* BL21(DE3). Large-scale (500 mL) LB Miller cultures were inoculated and grown at 37 °C shaking at 225 rpm until the optical density at 600 nm (OD_600nm_) reached 0.6 – 0.9, induced by addition of 1 mM isopropyl β- d-1-thiogalactopyranoside (IPTG), and grown for an additional 3 h under the same conditions. Cell pellets were harvested by centrifugation at 6,079 × *g* for 15 min at 4 °C and stored at −80 °C until resuspension for purification. Pellets were resdissolved in lysis buffer (20 mM Tris-HCl (pH 7.5), 350 mM NaCl 1 mM phenylmethylsulfonyl fluoride (PMSF), 10 mM imidazole, 5 mM β- mercaptoethanol (BME), and 1 mg/mL lysozyme) at 5 mL/g concentration and incubated at 4 °C for 30 min, followed by sonication (Fisherbrand Model 120; 2 – 4 30 s cycles at 60% intensity with bursts of 1 s on and 1 s off). The lysate was cleared by centrifugation at 18,138 × *g* for 1 h at 4 °C. Imidazole was added to a final concentration of 30 mM and then the cleared lysate was incubated with Ni-NTA agarose (Thermo Scientific HisPur resin; 2 mL bed volume) for 1 h at 4°C. After allowing the lysate to flow through, the column was washed sequentially with a high-salt buffer (20 mM Tris-HCl (pH 7.5), 500 mM NaCl, 5 mM BME, 1 mM PMSF, and 30 mM imidazole) and a low-salt buffer (20 mM Tris-HCl (pH 7.5), 100 mM NaCl, 5 mM BME, 1 mM PMSF, and 30 mM imidazole). Bound protein was eluted in stages with elution buffer (20 mM Tris-HCl (pH 7.5), 100 mM NaCl, 5 mM BME, and 150 mM imidazole). Fractions containing the protein were pooled and dialyzed overnight against dialysis buffer (20 mM Tris-HCl (pH 7.5), 100 mM NaCl, 5 mM BME, and 10% glycerol). The dialyzed solution was concentrated using a spin concentrator (Amicon Ultra-4 10,000 MWCO) by centrifugation at 4,000 × *g* at 4 °C. The protein concentration was quantified spectrophotometrically by measuring the absorbance at 280 nm (*A*_280nm_) using a microvolume spectrophotometer (DeNovix DS-11FX) and converting to concentration using extinction coefficients from the ExPASY ProtParam tool (ε_280nm_ = 15,930 M^−1^ cm^−1^ and 14,440 M^−1^ cm^−1^ for *E. coli* and *B. subtilis* DnaN, respectively). Protein purity was assessed by SDS-PAGE. Protein stocks were aliquoted, flash frozen in liquid nitrogen, and stored at −80 °C until use.

For validation, a separate batch of N-terminally His_6_-tagged *B. subtilis* was purchased from a commercial source (GenScript). Unless noted otherwise, binding assays were performed with this batch of protein.

### Peptide Synthesis

Commercially purchased solvents and reagents were used without further purification. Nα-Fmoc-protected amino acids and peptide synthesis reagents were purchased from Advanced ChemTech, ChemImpex International, Oakwood Chemical, and Gyros Protein Technologies. Peptides were synthesized manually or using a Gyros Protein Technologies PurePep^TM^ Chorus synthesizer. Peptides were synthesized following standard Fmoc solid-phase approaches on high-loading Rink MBHA resin (0.62 mmol/g resin). For solid-phase fluorescein labeling of the *E. coli*-derived peptides (QLSLPL and QADMA), 5 equivalents each of fluorescein isothiocyanate (FITC) and N,N’-diisopropylethylamine (DIEA) in N,N-dimethylformamide (DMF) were added to the peptide after removal of the final Fmoc protecting group. For solid-phase fluorescein labeling of the *B. subtilis-*derived peptides (QLSLF, QMGLF, and AMGLA), 5 equivalents each of 5(6)-carboxyfluorescein, ethyl cyanohydroxyiminoacetate (Oxyma), and N,N’-diisopropylcarbodiimide (DIC) were added to DMF for 20 minutes for pre-activation and then added to the peptide after removal of the final Fmoc protecting group. Peptides were globally deprotected and cleaved from resin by incubation with Reagent R (90% TFA, 5% thioanisole, 3% 1,2-ethanedithiol, and 2% anisole) for 2 h. After filtration through a cotton plug, rotary evaporation of TFA and precipitation with ice-cold diethyl ether yielded crude peptides. Peptides were purified on preparative C_18_ columns using reverse-phase high-performance liquid chromatography (RP-HPLC) on a Shimadzu Nexera HPLC system using gradients of water and acetonitrile (ACN) containing 0.1% trifluoroacetic acid (TFA). Peptide purity was evaluated by mass spectrometry (Table S1) and analytical HPLC (Figure S2) using a Luna 5 μm C18(2) column (150 × 4.6 mm) on an Agilent 1100 HPLC system with a flow rate of 0.3 mL/min (gradient: 5 – 95% solvent B over 30 minutes, solvent A = 0.1% TFA, B = 95% ACN, 5% water, 0.1% TFA).

### Fluorescence Polarization Binding Measurements

Binding affinities were determined using fluorescence polarization assays with fluorescein-tagged peptides and purified DnaN proteins. Proteins were thawed on ice and diluted in the same dialysis buffer used for purification, supplemented with 0.1% Pluronic acid F-68. Stock solutions of fluorescent peptides were prepared by dissolving peptides in 10 mM sodium phosphate buffer (pH 7.4) supplemented with 0.1% Pluronic acid F-68 to a concentration of 2 μM (100× the final assay concentration of 20 nM).

For each assay, a volume of 225 μL of protein was diluted in the supplemented dialysis buffer to the maximum concentration (10 μM for *E. coli* DnaN, 50 μM for *B. subtilis* DnaN) and used to prepare a 3-fold serial dilution (150 μL per dilution). For each protein dilution, a 1.5 μL volume of fluorescent peptide was added such that the final peptide concentration was 20 nM. Negative control samples were prepared by diluting 1.5 μL of fluorescent peptide into 150 μL of dialysis buffer supplemented with 0.1% Pluronic acid F-68. Each sample was then divided into 3 wells of a black 96-well plate (50 μL/well).

Data were collected on a BioTek Synergy H1M plate reader equipped with polarization filters (excitation: 485 nm/20 nm, emission: 528 nm/20 nm) using the fluorescence polarization program within the Gen5 software (version 3.08). Data were exported and fit to a 1:1 binding model accounting for the presence of both free and bound protein according to the following equation:

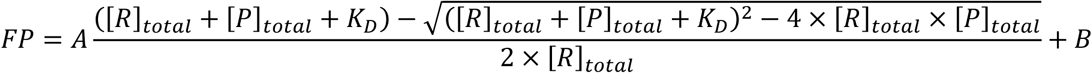

where A is the amplitude (maximum FP response), B is the baseline (minimum FP response), [*R*]_total_ is the peptide concentration (in μM), and [*P*]_total_ is the protein concentration (in μM).(61)

### Growth Curve Measurements

Liquid cultures were grown in LB Lennox and S7_50_-sorbitol media at 25 mL scale following the same approach reported previously for imaging culture growth.(35) OD_600nm_ was measured using a 1:1 dilution of growth curve sample in fresh media. Measurements were taken hourly until the cells entered early exponential phase, which took approximately two hours in LB Lennox media and three hours in S7_50_-sorbitol media, respectively. From then onward, OD_600nm_ was measured every half hour until the cells reached stationary phase. Growth rates and doubling times were determined via a linear fit to the exponential phase of the growth curves.

### Spot Dilution Growth Assays

Liquid cultures were grown in LB Lennox and S7_50_-sorbitol media at 25 mL scale following the same approach described for growth curve measurements. LB Lennox and S7_50_-sorbitol agar plates were freshly prepared with 1.5% agar on the day of spotting. When cultures reached OD_600nm_ ≈ 0.3, cell concentrations were normalized to OD_600nm_ = 0.2. These samples were then serially diluted 10-fold in media down to OD_600nm_ = 2 × 10^−6^. A 5 μL volume of each dilution was spotted on the agar plates. Images of the plates were recorded after approximately 16 h incubation at 37 °C.

### Marker Frequency Analysis and Whole-Genome Sequencing

Cultures were grown in S7_50_-sorbitol or LB Lennox media at 25 mL scale following the same approach described for growth curve measurements. When cultures reached OD_600nm_ ≈ 0.15, cells were harvested and concentrated by centrifugation in a series of steps at room temperature: first 10 min at 10,000 × *g*, then 5 min in a Fisher Scientific Centrific 225 Centrifuge on power level 8.5, and finally 3 min at 16,000 × *g* to remove all the supernatant. Cell pellets were flash frozen in liquid nitrogen and stored at −80 °C until use. Genomic DNA was extracted using the Qiagen DNeasy blood and tissue kit (catalog # 69504; Qiagen). DNA was sonicated using a Qsonica Q800L sonicator for 12 min at 20% amplitude to achieve an average fragment size of 170 bp. A DNA library was prepared using the NEBNext Ultra II kit (E7645; New England Biolabs) and sequenced using Illumina Nextseq2000. Sequencing reads were mapped to the *B. subtilis* PY79 genome (NCBI reference sequence NC_022898.1) using CLC Genomics Workbench (Qiagen). The mapped reads were normalized to the total number of reads and plotted in 10-kb bins using R.

### DNA Damage Survival Assays

Cell survival assays with 4-nitroquinoline 1-oxide (4-NQO) were performed as described previously.(35) In brief, exponentially growing cultures in 3 mL LB Lennox media were serially diluted and spread on LB Lennox agar plates containing different 4- NQO concentrations (0, 0.05, 0.075, or 0.1 μM). Plates were incubated overnight at 37 °C, colonies were enumerated, and the survival rate relative to the no drug plate was calculated. At least three independent experimental replicates were performed for each strain.

To assay survival to UV light, cells were cultured in LB Lennox media following the same approach as for 4-NQO survival assays. Cultures were serially diluted and spread on plain LB Lennox agar plates, then irradiated with different dosages (0, 10, 20, or 40 J/m^2^) of 254 nm UV light (Analytik Jena UVS-28 EL). Plates were incubated overnight, and survival rates were calculated as above. At least three independent experimental replicates were performed for each strain.

### Rifampicin Resistance Mutagenesis Assays

Spontaneous and UV-induced mutagenesis was assayed by measuring the rate of rifampicin resistance. Cultures were grown in LB Lennox media following a similar approach as for survival assays. Fresh cultures were prepared at 10 mL scale and incubated for 3.5 h, then pelleted by centrifugation in a Fisher Scientific Centrific Centrifuge on power level 8.5 for 5 min. Cell pellets were resuspended in 50 mL of 10 mM MgSO_4_. The resulting 50 mL solution was divided into two halves, one of which was not treated and one of which was irradiated with 254 nm UV light at a dose of 40 J/m^2^. Cells were pelleted by centrifugation in two stages, first at 10,000 × *g* for 6 min and then in a Fisher Scientific Centrific Centrifuge as on power level 8.5 for 5 min. Cell pellets were resuspended in 10 mL LB Lennox media and incubated overnight for 22 – 24 h at 37 °C shaking at 225 rpm. The following morning, LB Lennox agar plates were freshly prepared either with or without 10 μg/mL rifampicin (by a 1:1,000 dilution of a rifampicin stock in dimethylsulfoxide (DMSO)). Cultures were concentrated by centrifugation as described previously to OD_600nm_ ≈ 20 and spread on Rif^+^ plates. Concentrated cultures were serially diluted to OD_600nm_ ≈ 10^−5^ and spread on Rif^−^ plates. Colonies were enumerated after overnight incubation at 37 °C. Mutation rates were determined by normalizing the number of CFUs per mL on Rif^+^ plates by the number on Rif^−^ plates. At least five independent experimental replicates were performed for each strain.

### Cell Culture and Sample Preparation for Microscopy

Samples were prepared for microscopy exactly as described previously.(35) In brief, cultures were grown in S7_50_-sorbitol minimal medium until early exponential phase (OD_600nm_ ≈ 0.1 – 0.2), then aliquots were harvested and sparsely labeled with Janelia Fluor X 554 (JFX_554_) HaloTag ligand at 1 nM final concentration for Halo-DnaE and 500 pM final concentration for PolC-Halo for 15 min at 37 °C and shaking at 225 rpm. Samples were washed by resuspension in S7_50_-sorbitol media to remove free dye and concentrated by centrifugation, then deposited on pads prepared from 3% agarose (NuSieve) dissolved in media. Pads were sandwiched between cleaned coverslips and imaged immediately. *Microscopy:* Microscopy was performed using a custom fluorescence microscope described in detail previously.(35) In brief, an inverted microscope (Nikon Ti2-E) was equipped with a Nikon CFI Apo 100×/1.49 NA total internal reflection fluorescence (TIRF) objective lens, a Hamamatsu ImageEM C9100-23BKIT EMCCD camera, and Chroma filters. Highly inclined thin illumination,(62) or near-TIRF, excitation was provided by focusing 514 nm and 561 nm lasers (Coherent Sapphire) to the objective back focal plane (BFP) and translating the beams away from the BFP optical axis. Excitation sequences were automated with computer-controlled shutters (Vincent Uniblitz) and a computer-controlled stage (Mad City Labs) was used for translation between fields of view. White light transillumination was used to record brightfield images of cells.

A short integration time (13.9 ms) was used in all imaging experiments, allowing the detection of both static and mobile Halo-DnaE and PolC-Halo molecules. JFX_554_-labeled HaloTag fusions were imaged using a 561 nm laser power density of approximately 15 W/cm^2^ at the sample. To visualize DnaX-mYPet foci, 100 frames of 514 nm excitation were recorded at the start of each movie at approximately 1 W/cm^2^ power density. Imaging of PolC-mYPet foci used a 514 nm laser power density of approximately 5 mW/cm^2^.

### Image Analysis

Image analysis was performed using the MATLAB-based packages MicrobeTracker(63) (for segmentation of brightfield images) and u-track(64, 65) (for single-particle detection and tracking), in addition to custom MATLAB code. The analysis approach was described in detail previously and the same analysis parameters were used.(35) As described previously, the first 20 frames of 561 nm excitation were averaged for u-track analysis of DnaX-mYPet and PolC-mYPet foci.

### Data Analysis

Analysis of Halo-DnaE and PolC-Halo diffusion in short-exposure movies was performed exactly as described before by calculating an apparent diffusion coefficient *D*\* from the mean squared displacement (MSD);(35) as an alternative approach, diffusion was also analyzed with the analysis package Spot-On(48) with the same analysis parameters used previously.(35) Analysis of the average cellular localization of DnaX-mYPet and PolC-mYPet foci was likewise performed exactly following prior methods, as was radial distribution function analysis to quantify colocalization between Halo-DnaE or PolC-Halo molecules and DnaX-mYPet foci.(35)

### Imaging Dataset

Imaging datasets for main experimental results consisted of at least three biological replicates (separate imaging cultures) collected on at least two different days; for some control experiments, two replicates on separate days were collected. The number of imaging days, replicates, cells, and tracks or foci for all figures are listed in Table S14.

## Data, Materials, and Software Availability

The data and custom MATLAB analysis code from this study have been deposited in a Zenodo repository (DOI: 10.5281/zenodo.15001034). The Whole-Genome Sequencing results used for Marker Frequency Analysis were deposited to NCBI SRA with accession number PRJNA1232650.

## Acknowledgments

We thank Thrall lab members for assistance and helpful discussions, Xheni Karaboja for technical assistance, and the Indiana University Center for Genomics and Bioinformatics for high throughput sequencing. We thank Joseph Loparo (Harvard Medical School) for sharing bacterial strains and Luke Lavis (Howard Hughes Medical Institute Janelia Research Campus) for providing Janelia Fluor dyes. Support for this work comes from National Institutes of Health R01GM141242 (to X.W.), R01GM143182 (to X.W.), R01AI172822 (to X.W.), and R15GM151677 (to E.S.T.). This research is a contribution of the GEMS Biology Integration Institute, funded by the National Science Foundation DBI Biology Integration Institutes Program, Award Number 2022049 (to X.W.). Additional support comes from the Camille and Henry Dreyfus Foundation Jean Dreyfus Lectureship under Award Number BL-22-012 (to L.G.O. and E.S.T), the Fordham College at Rose Hill Undergraduate Research Grant program (to L.G.O., M.N.D., A.F.C., A.P.C., and S.H.), the Fordham University Clare Boothe Luce program (to A.F.C. and E.E.H.), and the Fordham University Faculty Research Grant program (to N.S. and E.S.T.).

## Author Contributions

E.S.T., X.W., and N.S. designed research; L.G.O., M.N.D., N.K.L., A.F.C., A.P.C., L.E.W., S.H., E.E.H., S.J.R., N.S., and E.S.T. performed research; M.N.D., N.K.L, A.F.C., L.E.W., S.J.R., N.S., X.W., and E.S.T. contributed new reagents; L.G.O., M.N.D., N.K.L., A.F.C., A.P.C., S.H., E.E.H., N.S., X.W., and E.S.T. analyzed data; and L.G.O., N.K.L., N.S., X.W., and E.S.T. wrote the paper.

