## Supplementary Information for "“The *B. subtilis* replicative polymerases bind the sliding clamp with different strengths to tune replication processivity and fidelity”"

**This PDF file includes:**

Figures S1 to S8
Tables S1 to S14
Supplementary Methods
Supplementary References

### Supplementary Figures

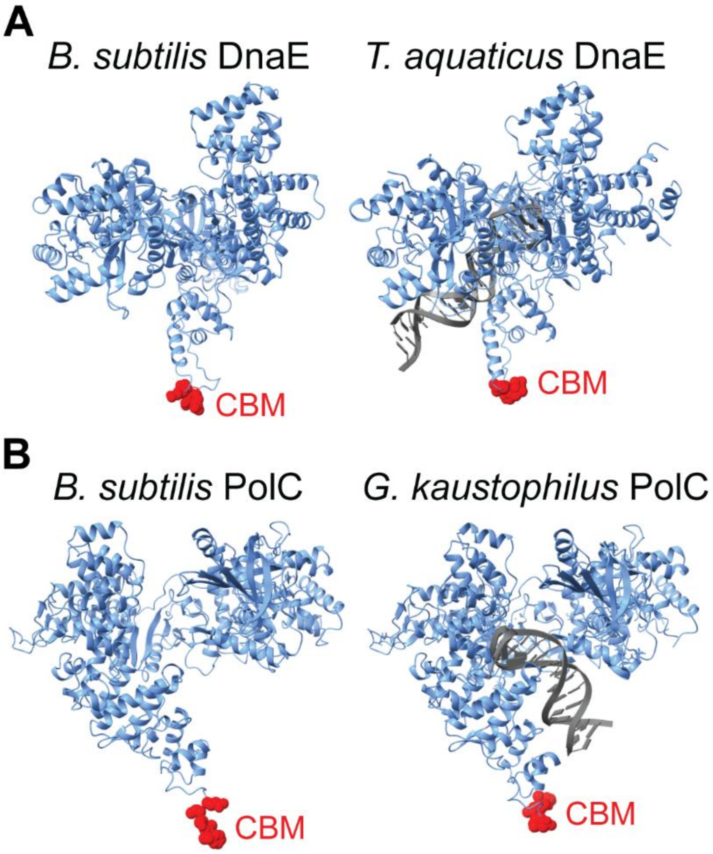


**Figure S1.** *B. subtilis* polymerase structures and comparisons to homologs. (A) AlphaFold predicted structure of *B. subtilis* DnaE (AlphaFold DB: AF-O34623-F1) (left) and experimental crystal structure of *T. aquaticus* DnaE (PDB: 3E0D)(1) (right). (B) AlphaFold predicted structure of *B. subtilis* PolC (AlphaFold DB: AF-P13267-F1) (left) and experimental crystal structure of *G. kaustophilus* PolC (PDB: 3F2B)(2) (right). The experimental PolC structure contains truncations of the N-terminal domain (residues 1 – 232) and exonuclease domain (residues 413 – 616). The corresponding truncations were made to the AlphaFold structure for comparison. CBMs are highlighted in red.

| CBM | Analytical HPLC Chromatogram |
|---|---|
| **QLSLPL** | 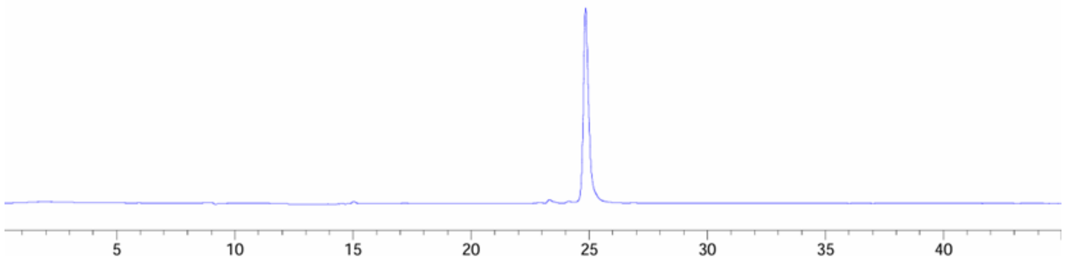 |
| **QADMA** | 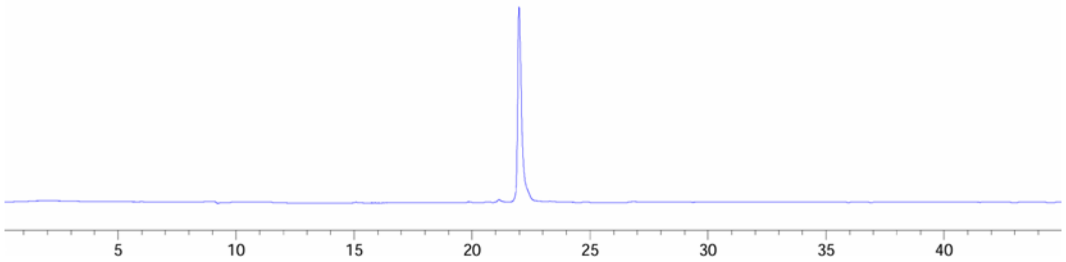 |
| **QLSLF (PolC context)** | 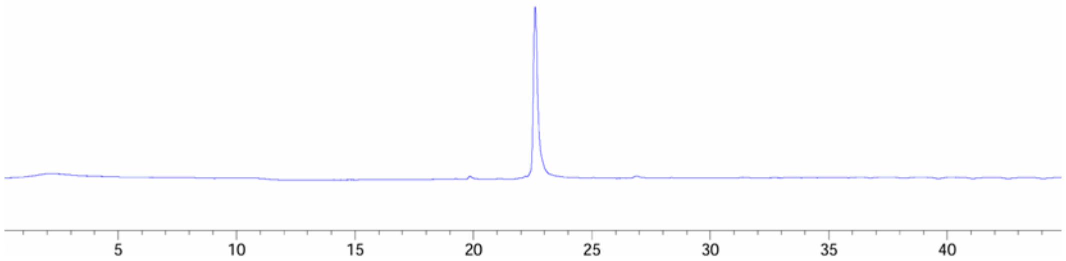 |
| **QMGLF** | 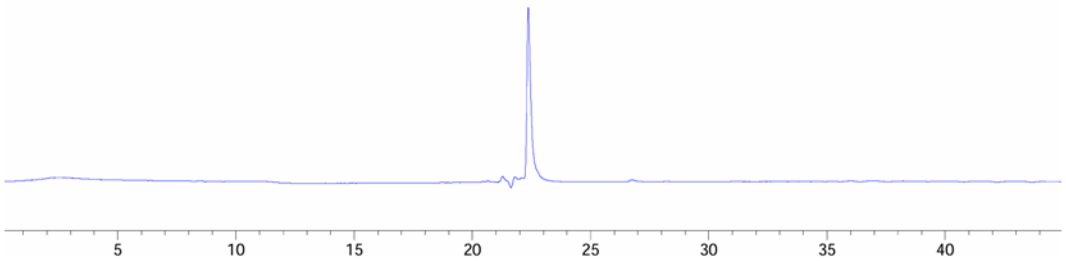 |
| **AMGLA** | 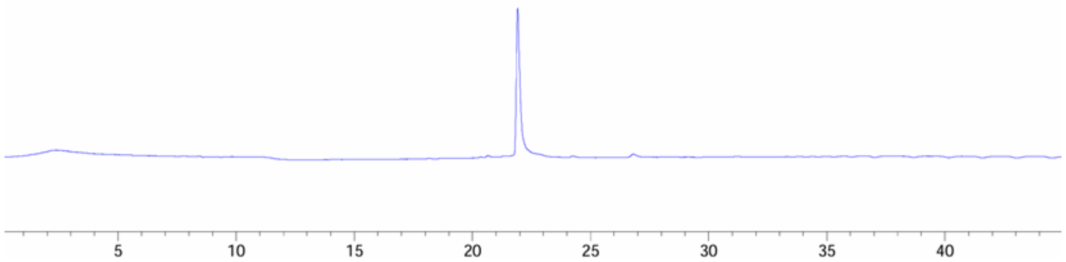 |
| **QLSLF (DnaE context)** | 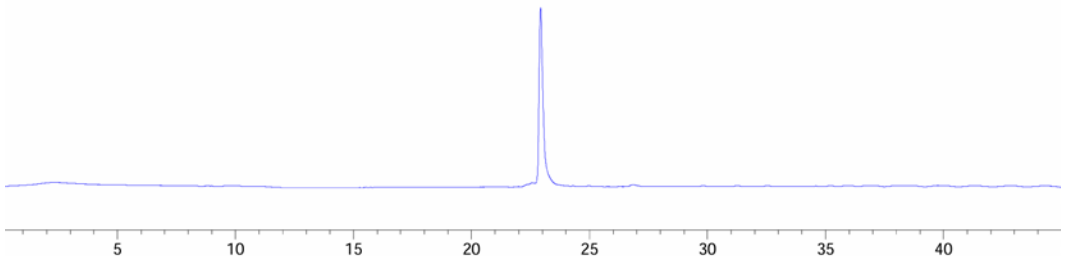 |

**Figure S2.** Analytical HPLC chromatograms for all peptides.

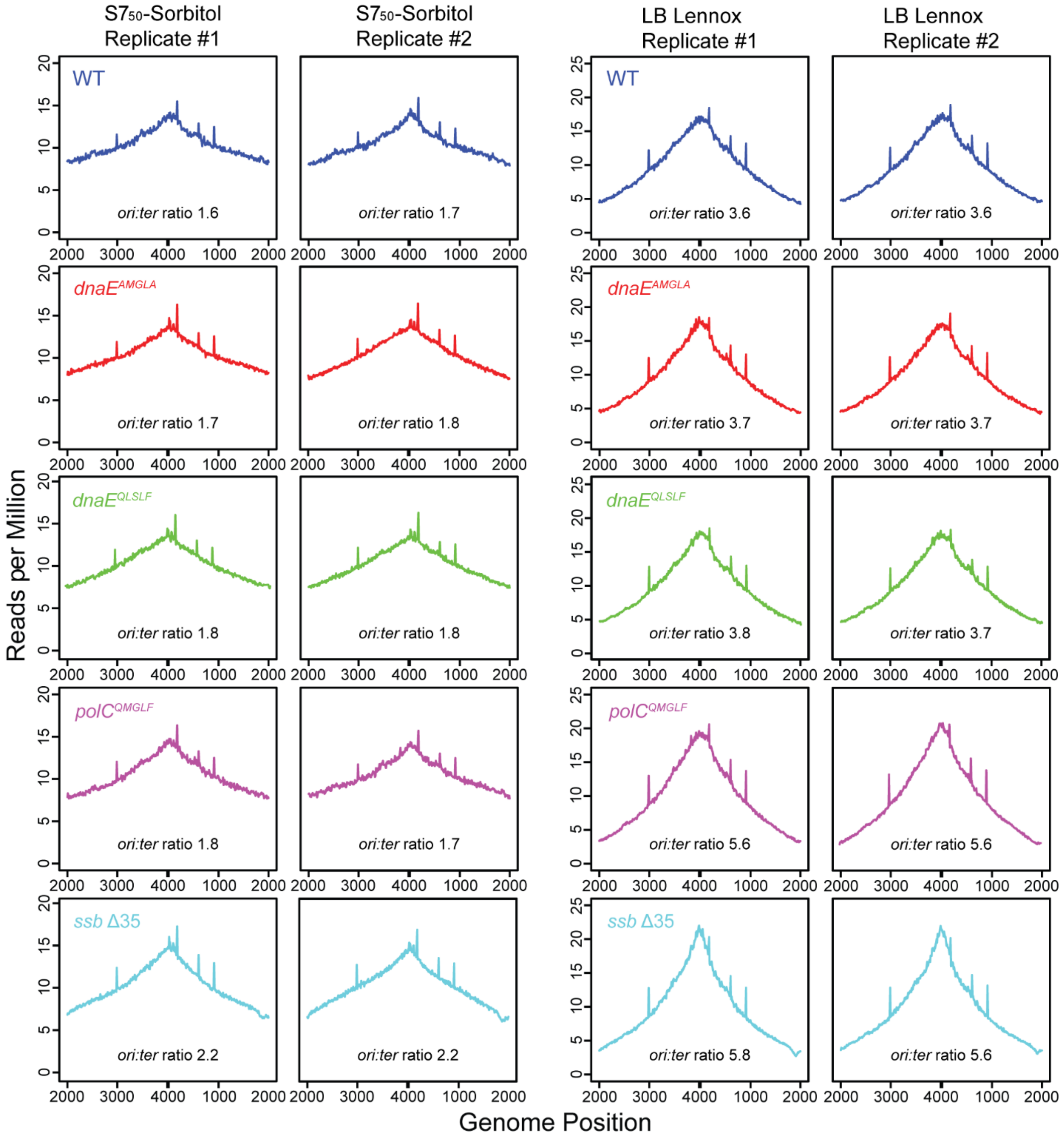


**Figure S3.** Whole-genome sequencing analysis for WT and CBM mutant strains of DnaE and PolC. Individual plots showing the normalized number of reads vs. genome location for (top to bottom) WT, *dnaE*$^{AMGLA}$, *dnaE*$^{QLSLF}$, *polC*$^{QMGLF}$, and *ssb* Δ35 strains in (left) S7$_{50}$-sorbitol and (right) LB Lennox media, respectively.

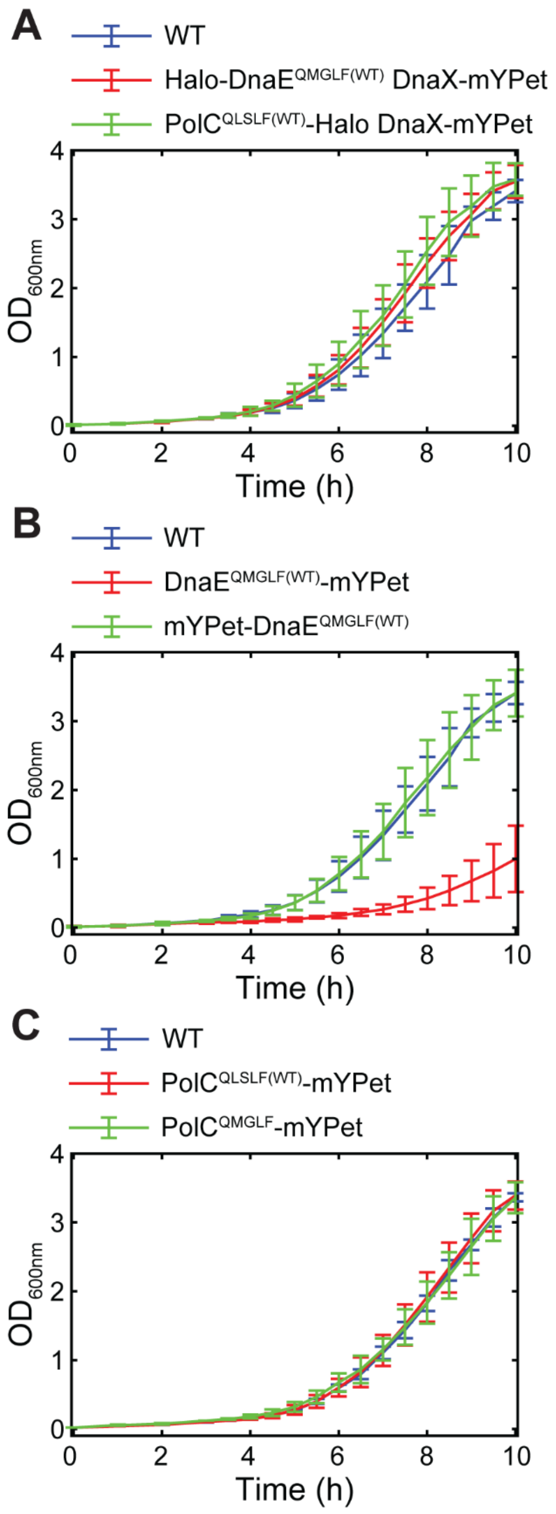


**Figure S4.** Liquid culture growth curves for imaging strains in S7$_{50}$-sorbitol media. (A) WT, Halo-DnaE$^{QMGLF(WT)}$ DnaX-mYPet, and PolC$^{QLSLF(WT)}$-Halo DnaX-mYPet. (B) WT, DnaE$^{QMGLF(WT)}$-mYPet, and mYPet-DnaE$^{QMGLF(WT)}$. (C) WT, PolC$^{QLSLF(WT)}$-mYPet, and PolC$^{QMGLF}$-mYPet.

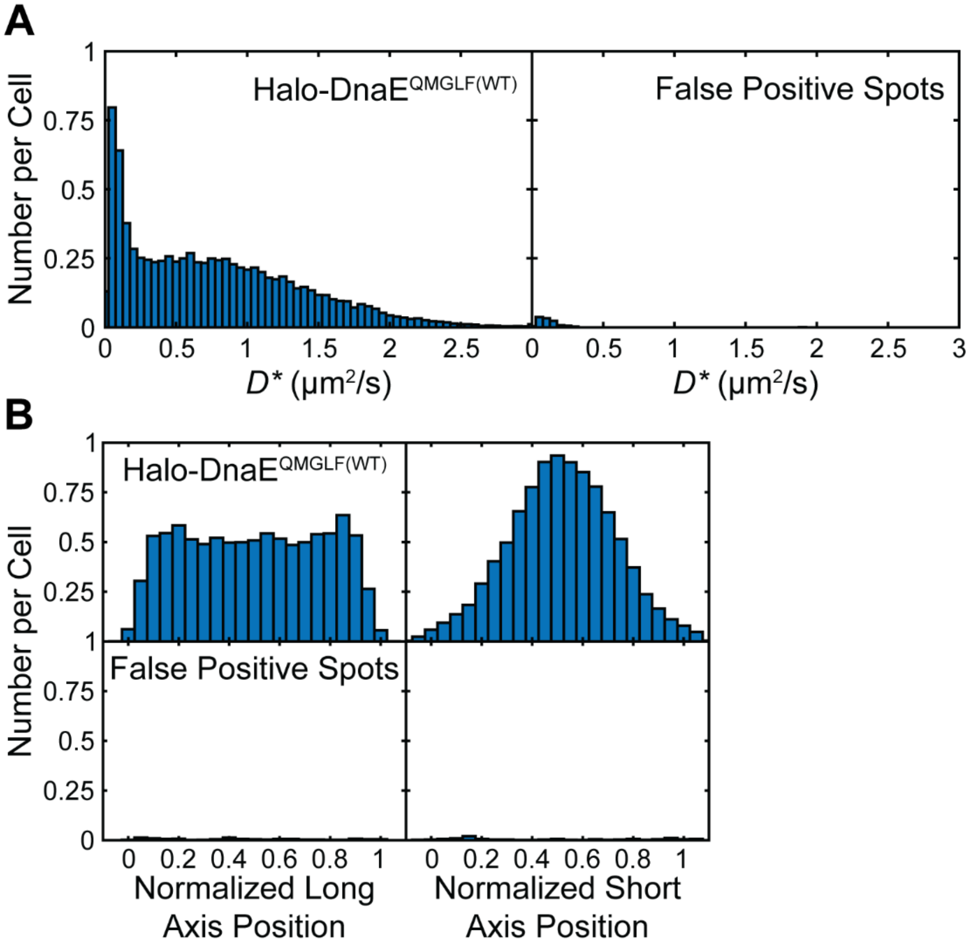


**Figure S5.** Comparison of false positive signal to Halo-DnaE signal. (A) Apparent diffusion coefficient ($D^*$) distributions for (left) Halo-DnaE$^{QMGLF(WT)}$ and (right) false positive spots on a per cell basis. (B) Average cellular localization projected along normalized long and short cell axes for (left) Halo-DnaE$^{QMGLF(WT)}$ and (right) false positive spots on a per cell basis.

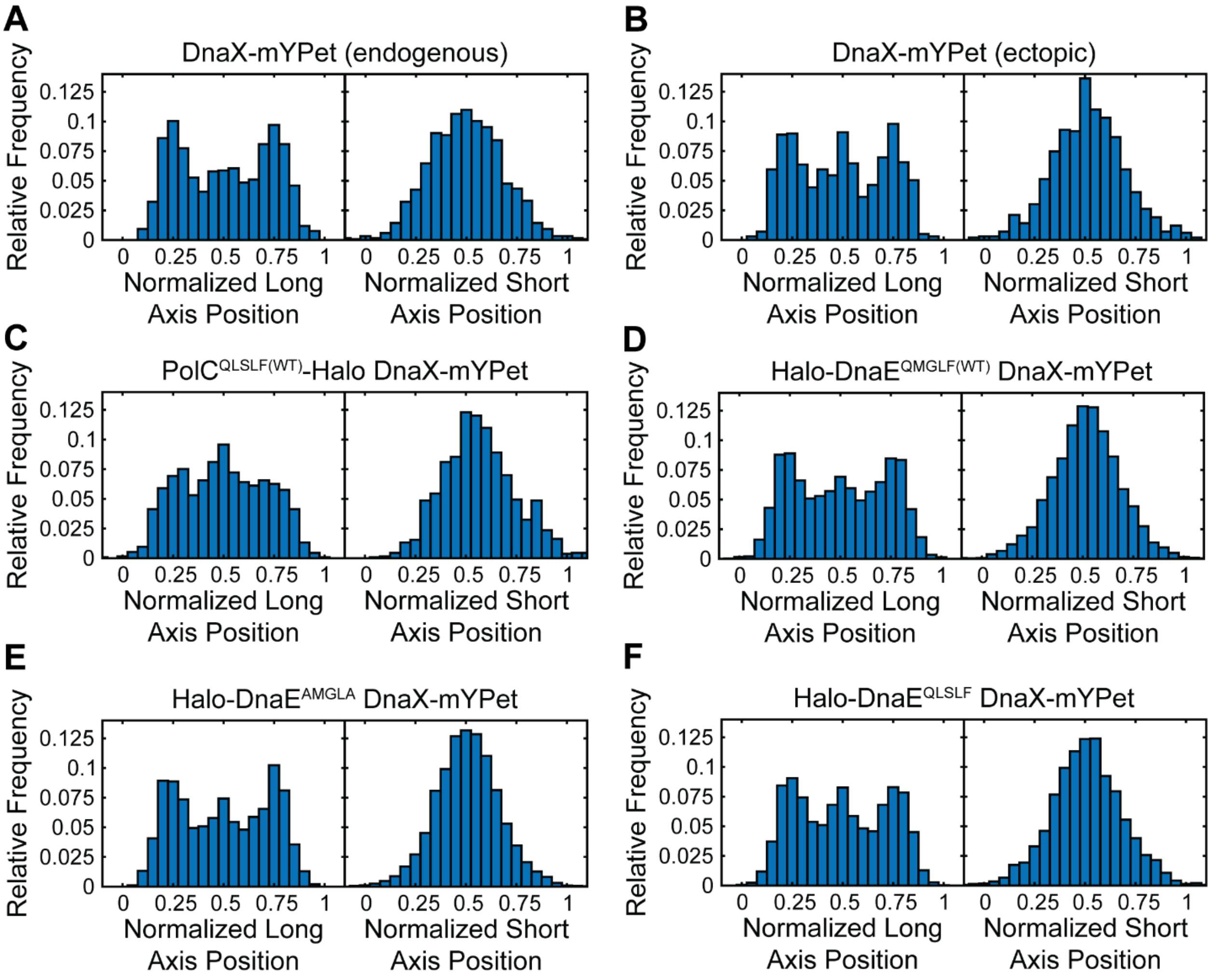


**Figure S6.** Average cellular localization of DnaX-mYPet foci projected along normalized long and short cell axes. (A) Endogenous DnaX-mYPet. (B) Ectopic DnaX-mYPet. (C) PolC-Halo DnaX-mYPet. (D) Halo-DnaE$^{QMGLF(WT)}$ DnaX-mYPet. (E) Halo-DnaE$^{AMGLA}$ DnaX-mYPet. (F) Halo-DnaE$^{QLSLF}$ DnaX-mYPet.

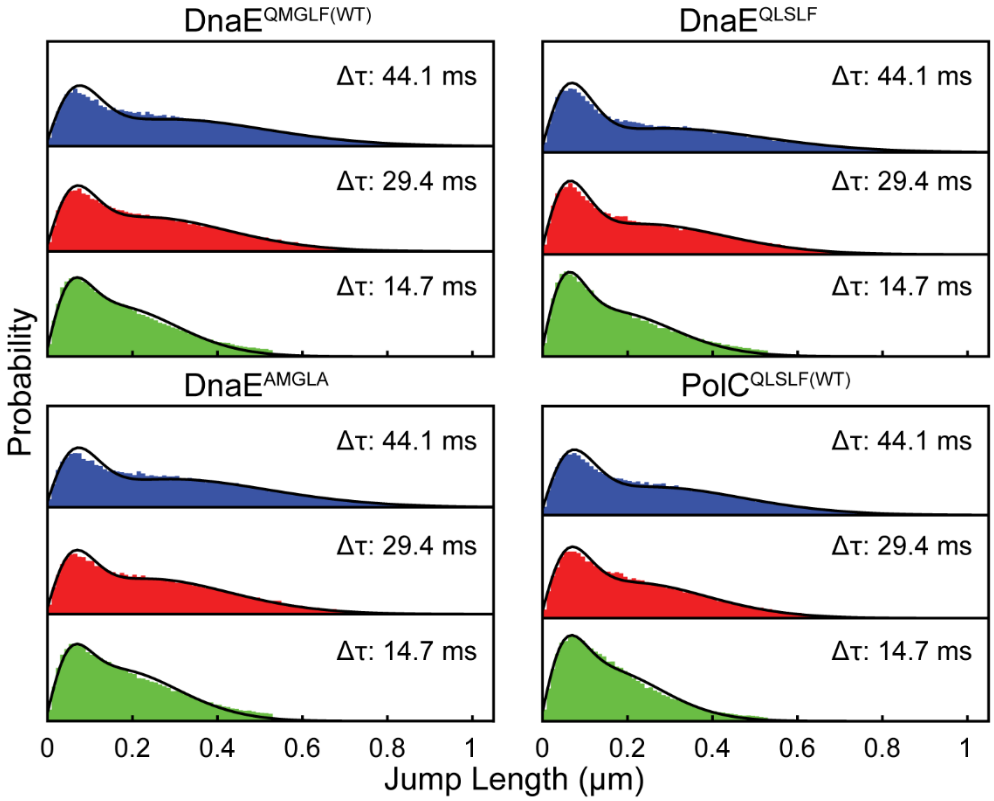


**Figure S7.** Spot-On diffusion analysis for Halo-DnaE and PolC-Halo, showing jump length distributions and corresponding two-population fits. (Top left) Halo-DnaE$^{QMGLF(WT)}$, (bottom left) Halo-DnaE$^{AMGLA}$, (top right) Halo-DnaE$^{QLSLF}$, and (bottom right) PolC$^{QLSLF(WT)}$-Halo.

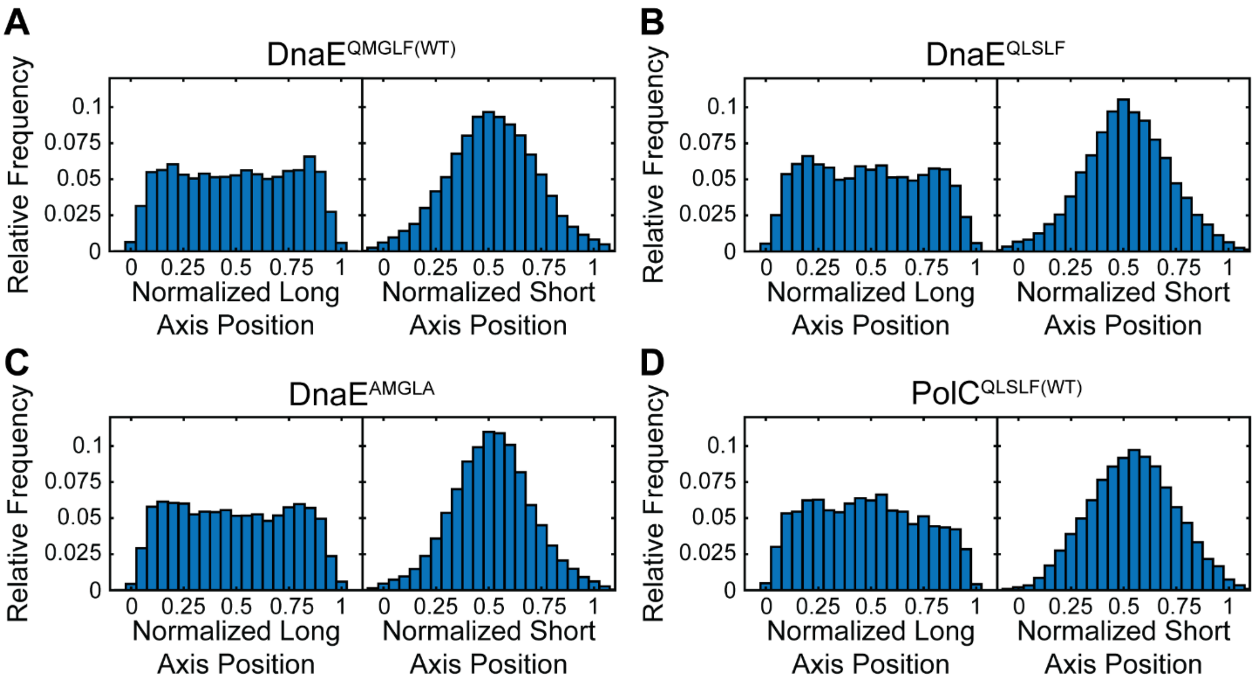


**Figure S8.** Average cellular localization of single Halo-DnaE or PolC-Halo molecules projected along normalized long and short cell axes. (A) Halo-DnaE$^{QMGLF(WT)}$. (B) Halo-DnaE$^{QLSLF}$. (C) Halo-DnaE$^{AMGLA}$. (D) PolC$^{QLSLF(WT)}$-Halo.

### Supplementary Tables

**Table S1.** MALDI Mass Spectrometry Characterization of CBM Peptides. FITC = fluorescein-5-thiocarbamoyl, FAM = fluorescein-5(6)-carbonyl, Ahx = 6-aminohexanoyl; -OH indicates a C-terminal carboxylic acid; $-NH_2$ indicates a C-terminal amide.

| **CBM** | **Peptide Name** | **Full Sequence** | **Expected *m/z* (M+H$^+$)** | **Observed *m/z* (M+H$^+$)** |
|---|---|---|---|---|
| QLSLPL | NSC1,1 | FITC-Ahx-PAQLSLPLYL-$NH_2$ | 1615.8 | 1615.9 |
| QADMA | NSC2,1 | FITC-Ahx-IGQADMAGV-$NH_2$ | 1362.5 | 1362.1 |
| QLSLF (PolC context) | PolC | FAM-Ahx-DQNQLSLF-OH | 1435.6 | 1435.9 |
| QMGLF | AC10B | FAM-Ahx-DDQMGLFLDE-$NH_2$ | 1652.6 | 1653.0 |
| AMGLA | AC11B | FAM-Ahx-DDAMGLALDE-$NH_2$ | 1519.6 | 1520.1 |
| QLSLF (DnaE context) | AC12B | FAM-Ahx-DDQLSLFLDE-$NH_2$ | 1664.7 | 1665.0 |

**Table S2.** Doubling times (mean ± standard deviation) for WT and PolC and DnaE CBM mutant strains for growth shaking at 225 rpm at 37 °C in minimal $S7_{50}$-sorbitol media and rich LB Lennox media. (Note: * indicates statistically significant difference at the $p < 0.05$ level relative to WT for same growth condition.)

| **Strain / Media** | **WT** | **DnaE-AMGLA** | **DnaE-QLSLF** | **PolC-QMGLF** |
|---|---|---|---|---|
| $S7_{50}$-Sorbitol (min) | 60 ± 1 | 58 ± 1 | 59.1 ± 0.6 | 58 ± 2 |
| LB Lennox (min) | 31 ± 1 | 28 ± 2 * | 30 ± 1 | 41 ± 3 * |

**Table S3.** Marker frequency analysis (MFA) of average *ori:ter* ratio (mean ± standard deviation) for WT and PolC and DnaE CBM mutant strains for growth shaking at 225 rpm at 37 °C in minimal $S7_{50}$-sorbitol media and rich LB Lennox media.

| **Strain / Media** | **WT** | ***dnaE$^{AMGLA}$*** | ***dnaE$^{QLSLF}$*** | ***polC$^{QMGLF}$*** | ***ssb D35*** |
|---|---|---|---|---|---|
| $S7_{50}$-Sorbitol | 1.65 ± 0.07 | 1.75 ± 0.07 | 1.80 ± 0.00 | 1.75 ± 0.07 | 2.20 ± 0.00 |
| LB Lennox | 3.60 ± 0.00 | 3.70 ± 0.00 | 3.75 ± 0.07 | 6.1 ± 0.7 | 5.7 ± 0.1 |

**Table S4.** Mutagenesis for different strains in untreated cells or cells treated with 40 J/m$^2$ UV light measured by the rate of rifampicin resistance (Rif$^R$) (mean ± standard deviation). (Note: * indicates statistically significant difference at the $p < 0.05$ level relative to WT for same treatment condition. All differences between untreated and treated for the same strain are statistically significant at the $p < 0.05$ level.)

| **Strain** | **WT** | | ***dnaE*$^{AMGLA}$** | | ***dnaE*$^{QLSLF}$** | |
|---|---|---|---|---|---|---|
| **Condition** | **Untreated** | **Treated** | **Untreated** | **Treated** | **Untreated** | **Treated** |
| **Rif$^R$ (per 10$^8$)** | 0.7 ± 0.2 | 15 ± 5 | 0.9 ± 0.8 | 22 ± 10 | 1.1 ± 0.5 | 43 ± 5 * |

**Table S5.** Doubling times (mean ± standard deviation) for WT and imaging strains for growth shaking at 225 rpm at 37 °C in minimal S7$_{50}$-sorbitol media. (Note: * indicates statistically significant difference at the $p < 0.05$ level relative to WT for same growth condition.)

| **Strain** | **Doubling Time (min)** | **Strain** | **Doubling Time (min)** |
|---|---|---|---|
| WT | 59 ± 2 | mYPet-DnaE$^{QMGLF(WT)}$ | 101 ± 23 * |
| Halo-DnaE$^{QMGLF(WT)}$<br>DnaX-mYPet | 58 ± 3 | DnaE$^{QMGLF(WT)}$-mYPet | 57 ± 4 |
| PolC$^{QLSLF(WT)}$-Halo<br>DnaX-mYPet | 58 ± 4 | PolC$^{QLSLF(WT)}$-mYPet | 59 ± 2 |
| | | PolC$^{QMGLF}$-mYPet | 61 ± 1 |

**Table S6.** Fold-change in number of colony-forming units (CFUs) per mL (mean ± standard deviation) in JFX$_{554}$-labeled vs. mock-labeled cultures after 1 h incubation. (JFX$_{554}$ concentrations: 1 nM for Halo-DnaE, 500 pM for PolC-Halo).

| **Strain** | **Fold-Change in CFUs/mL** | |
|---|---|---|
| | **Mock-Labeled** | **Labeled** |
| Halo-DnaE DnaX-mYPet | 2.6 ± 0.5 | 2.7 ± 0.7 |
| PolC-Halo DnaX-mYPet | 2.1 ± 0.2 | 2.3 ± 0.5 |

**Table S7:** Cell length and width and number of DnaX or PolC foci per cell (mean ± standard error of the mean (S.E.M.)) (Note: ‡ indicates difference was not statistically significant at the $p < 0.05$ level. For DnaE and PolC HaloTag fusions, the comparison is made to the corresponding strain with the DnaX-mYPet fusion alone; for PolC$^{QMGLF}$-mYPet, the comparison is made to PolC$^{QLSLF(WT)}$. For DnaX-mYPet (ectopic), the comparison is made to DnaX-mYPet (endogenous).)

| **Strain** | **Cell Length (mm)** | **Cell Width (mm)** | **Number of Foci per Cell** |
|---|---|---|---|
| DnaX-mYPet (endogenous) | 3.09 ± 0.03 | 0.641 ± 0.003 | 1.62 ± 0.03 |
| DnaX-mYPet (ectopic) | 2.95 ± 0.03 | 0.589 ± 0.002 | 1.39 ± 0.03 |
| Halo-DnaE$^{QMGLF(WT)}$ DnaX-mYPet | 3.04 ± 0.02 | 0.665 ± 0.001 | 1.60 ± 0.02 ‡ |
| Halo-DnaE$^{AMGLA}$ DnaX-mYPet | 3.22 ± 0.02 | 0.686 ± 0.002 | 1.71 ± 0.02 |
| Halo-DnaE$^{QLSLF}$ DnaX-mYPet | 3.17 ± 0.02 | 0.650 ± 0.002 | 1.56 ± 0.02 |
| PolC$^{QLSLF(WT)}$-Halo DnaX-mYPet | 3.75 ± 0.03 | 0.636 ± 0.002 | 1.42 ± 0.03 ‡ |
| PolC$^{QLSLF(WT)}$-mYPet | 3.58 ± 0.04 | 0.641 ± 0.003 | 1.71 ± 0.04 |
| PolC$^{QMGLF}$-mYPet | 4.36 ± 0.04 | 0.611 ± 0.003 | 1.78 ± 0.04 ‡ |

**Table S8:** Halo-DnaE and PolC-Halo diffusion coefficient distribution fit parameters from MSD analysis (± uncertainties from 95% fit confidence intervals).

| **Strain** | $D_1$ **(μm²/s)** | $A_1$ | $D_2$ **(μm²/s)** | $A_2$ | $D_3$ **(μm²/s)** | $A_3$ |
|---|---|---|---|---|---|---|
| Halo-DnaE$^{QMGLF(WT)}$ | 0.084 ± 0.004 | 0.174 ± 0.011 | 0.306 ± 0.035 | 0.161 ± 0.020 | 1.0981 ± 0.040 | 0.666 ± 0.031 |
| Halo-DnaE$^{AMGLA}$ | 0.082 ± 0.004 | 0.162 ± 0.011 | 0.335 ± 0.048 | 0.149 ± 0.025 | 1.125 ± 0.048 | 0.689 ± 0.036 |
| Halo-DnaE$^{QLSLF}$ | 0.077 ± 0.005 | 0.205 ± 0.019 | 0.256 ± 0.040 | 0.157 ± 0.024 | 1.081 ± 0.055 | 0.637 ± 0.043 |
| PolC$^{QLSLF(WT)}$-Halo | 0.084 ± 0.004 | 0.186 ± 0.012 | 0.294 ± 0.034 | 0.190 ± 0.025 | 0.878 ± 0.036 | 0.624 ± 0.037 |

**Table S9:** Halo-DnaE and PolC-Halo diffusion coefficients and populations from Spot-On analysis.

| **Strain** | $D_{static}$ **(μm²/s)** | $A_{\mathrm{static}}$ | $D_{\mathrm{free}}$ **(μm²/s)** | $A_{\mathrm{free}}$ |
|---|---|---|---|---|
| Halo-DnaE$^{QMGLF(WT)}$ | 0.029 | 0.217 | 0.959 | 0.783 |
| Halo-DnaE$^{AMGLA}$ | 0.025 | 0.198 | 0.982 | 0.802 |
| Halo-DnaE$^{QLSLF}$ | 0.022 | 0.261 | 0.975 | 0.739 |
| PolC$^{QLSLF(WT)}$-Halo | 0.027 | 0.243 | 0.800 | 0.757 |

**Table S10:** Value of the mean radial distribution function $g(r)$ for DnaE-DnaX or PolC-DnaX colocalization at the second smallest value of $r$ (the maximum of the $g(r)$ curve) and the standard error of the mean (S.E.M.) at that $r$ value for the 100 calculated $g(r)$ curves.

| Strain | $g(r)$ ± S.E.M. |
|---|---|
| Halo-DnaE$^{QMGLF(WT)}$ | 1.272 ± 0.005 |
| PolC$^{QLSLF(WT)}$-Halo | 2.82 ± 0.02 |
| Halo-DnaE$^{AMGLA}$ | 1.633 ± 0.007 |
| Halo-DnaE$^{QLSLF}$ | 2.91 ± 0.02 |

**Table S11:** Oligonucleotides used in this study. (Lowercase letters indicate bases that do not prime on the template but provide homology for Gibson assembly of PCR fragments.)

| Number | Designation | Sequence (5′-3′) |
|---|---|---|
| oEST082 | Ec-pET28b-his6-dnaN-for | ggcagcagccaccatcaccaccatcatagcagcggcATGAAATTTACCGTAGAACG |
| oEST083 | Ec-pET28b-dnaN-rev | gctttgttagcagccggatcTTACAGTCTCATTGGCATGA |
| oEST084 | Bs-pET28b-his6-dnaN-for | ggcagcagccaccatcaccaccatcatagcagcggcATGAAATTCACGATTCAAAA |
| oEST085 | Bs-pET28b-dnaN-rev | gctttgttagcagccggatcTTAATAGGTTCTGACAGGAA |
| oEST086 | pET28b-his6-for | GATCCGGCTGCTAACAAAGC |
| oEST087 | pET28b-his6-rev | tggtgatggtggctgctgccCATGGTATATCTCCTTCTTAAAG |
| oEST118 | yvbJ-upstream-for | CTTCCTGTACCGATGATGAACATC |
| oEST119 | yvbJ-Nter-rev | TTTAAGTCATAATGGTCAAATACCAGC |
| oEST120 | yvbJ-Nter-cat-iso-for | tttgaccattatgacttaaaCCATACGGCAATAGTTACCCTTAT |
| oEST121 | dnaX-yvbJ-Cter-iso-rev | gaaaatccactttttctgttGAATCGACAGAAGGTGCACGGTAA |
| oEST122 | yvbJ-Cter-for | AACAGAAAAAGTGGATTTTCGAAG |
| oEST123 | yvbJ-downstream-rev | AATAGGAAGGACGTATACAGATGT |
| oML83 | sequencing oligo in *kan* | CCTCATCCTCTTCATCCTC |
| oWX225 | region at C-ter of *polC* | GACTTGATGCTTCACCGAAGG |
| oWX228 | region at C-ter of *polC* | CGCTCTAGAGCTTACTTTGGCACATTCCTC |
| oWX323 | region downstream of *polC* | GGTCCCTGTCCTGATCCCTCGAGGAACAGTGACAATTGGTTTTG |
| oWX324 | region downstream of *polC* | CGCGGGATCGAGATCTGCATTAATAATATGGAAAGCAGAATTTCTC |
| oWX438 | universal oligo for *loxP* cassettes | GACCAGGGAGCACTGGTCAAC |
| oWX439 | universal oligo for *loxP* cassettes | TCCTTCTGCTCCCTCGCTCAG |
| oWX900 | region upstream of *dnaE* | TCTCGTATTGCCTTCGAGTATGCCGCG |
| oWX902 | sequencing *dnaE* | GCGGACCTATTGAGAAGGGCGGTCAGC |

| | | |
|---|---|---|
| oWX903 | region upstream of *dnaE* | CTGAGCGAGGGAGCAGAAGGATCCGTATAACAAAAAGCGA ATCGTTACC |
| oWX904 | region downstream of *dnaE* | GGTAGTTGACCAGTGCTCCCTGGTCAATTGGAGCAATTAT GTGATCCTG |
| oWX905 | region downstream of *dnaE* | TCACCTCCTGTTTGATGCATATCCAGC TCACCTCCTGTTTGATGCATATCCAGC |
| oWX907 | sequencing *dnaE* | AGTTACAACTGCTACCATGTTCCC |
| oWX916 | region upstream of *polC* gene | CAAAAGCTTGCGGAAGACCAAGAGCGG |
| oWX919 | region inside of *polC* gene | GAGCAGAAGGATCCCTGAGAAATTCTGCTTTCCATATTAG |
| oWX920 | region downstream *polC spec* and region downstream *polC mYpet cat* | TGACCAGTGCTCCCTGGTCAAATTCTGCTTCTATGCATAC ATAAGCG |
| oWX921 | region downstream *polC spec* and region downstream *polC mYpet cat* | CTGATCAGCTTATTCGCAAAACCTTGG |
| oWX2235 | region upstream of *ssb* | CTGATAATGGAATGGGAAGTACGC |
| oWX2236 | region upstream of *ssb* | CTGAGCGAGGGAGCAGAAGGATTAGAATGGAAGATCATCA TCCGAG |
| oWX2238 | for *ssb* Δ35 | CTGAGCGAGGGAGCAGAAGGATTACCCAAATGGATTATCA TTTTGGCC |
| oWX2239 | region downstream of *ssb* | GTTGACCAGTGCTCCCTGGTCTGTGATTATCGCCTAAAAT GAAAAAG |
| oWX2240 | region downstream of *ssb* | ATGATACAAATCATTCGCTTCGCC |
| oWX2241 | for *dnaE* Q885A | GATTCATCTAAAAACAATCCCATCGCGTCATCGTCCGCGG CGAATAGC |
| oWX2242 | for *dnaE* Q885A | GCTATTCGCCGCGGACGATGACGCGATGGGATTGTTTTTA GATGAATC |
| oWX2249 | for *dnaE* Q885A | GTCATCGTCCGCGGCGAATAGC |
| oWX2354 | for *dnaE* | CAATCCCATCGCGTCATCGTCCGC |
| oWX2355 | for *dnaE* Q885A F889A | GCGGACGATGACGCGATGGGATTGGCTTTAGATGAATCGT TTTCAATTAAGC |
| oWX2357 | for *dnaE* M886L G887S | GCTATTCGCCGCGGACGATGACCAACTGTCATTGTTTTTA GATGAATCGTTTTC |
| oWX3633 | region upstream of *polC* gene | GAACAATCCCATTTGGTTTTGATCAGGCAGTGAC |
| oWX3634 | for *polC* CBM mutant | AAAACCAAATGGGATTGTTCTAATATGGAAAGCAGAATTT CTC |
| oWX3692 | region downstream *polC mYpet* | AAAACCAAATGGGATTGTTCCTCGAGGGATCAGGACAGGG ACC |

**Table S12:** Plasmids used in this study.

| Number | Designation | Containing Strain | Source or Reference |
|---|---|---|---|
| pEST005 | *pET-28b-mCherry-his*$_6$ | EST149 | Gift of Joseph Loparo (Harvard Medical School); plasmid pET012 |
| pEST007 | *pET-28b-his*$_6$*-Ec-dnaN* | EST199 | This study |
| pEST009 | *pET-28b-his*$_6$*-Bs-dnaN* | EST201 | This study |
| pWX318 | *mYpet b.s. cat* | cWX514 | (3) |
| pWX340 | *dnaX-mYpet b.s.Ω cat* | cWX571 | (3) |
| pWX466 | *loxP-spec-loxP* | cWX856 | (4) |
| pWX470 | *loxP-kan-loxP* | cWX860 | (4) |

**Table S13:** *B. subtilis* bacterial strains used in this study.

| Number | Designation or description | Relevant genotype | Construction or source strain designation | Reference |
|---|---|---|---|---|
| BWX2231 | PolC spec | *PY79 polC loxP spec* | Transformation: *polC loxP spec* → EST003 | This study |
| EST001 | *E. coli* cloning strain | DH5a | — | — |
| EST003 | *B. subtilis* prototrophic wild-type strain | PY79 | — | (5, 6) |
| EST007 (BWX5094) | SSB Δ35 | PY79 *ssb Δ35 loxP kan* | Transformation: *ssb* D*35 loxP kan* → EST003 | This study |
| EST009 (BWX5096) | DnaE loxP kan | PY79 *dnaE loxP kan* | Transformation: *dnaE loxP kan* → EST003 | This study |
| EST013 (BWX5106) | DnaE$^{AMGLF}$ | PY79 *dnaE*$^{AMGLF}$ *loxP kan* | Transformation: *dnaE*$^{AMGLF}$ *loxP kan* → EST003 | This study |
| EST043 (BWX5217) | DnaE$^{AMGLA}$ | PY79 *dnaE*$^{AMGLA}$ *loxP kan* | Transformation: *dnaE*$^{AMGLA}$ *loxP kan* → EST003 | This study |
| EST047 (BWX5220) | DnaE$^{QLSLF}$ | PY79 *dnaE*$^{QLSLF}$ *loxP kan* | Transformation: *dnaE*$^{QLSLF}$ *loxP kan* → EST003 | This study |
| EST051 | DnaE-mYPet | PY79 *dnaE-mYpet cat* | Gift of Joseph Loparo (Harvard Medical School); strain ET017 | — |
| EST053 (BWX519) | DnaX-mYPet | PY79 *dnaX-mYpet cat Ω pWX340a* | Strain BWX519 from plasmid pWX340 | (3) |
| EST057 (BWX499) | PolC-mYPet | PY79 *polC-mYPet cat* | Transformation: *polC-mYPet cat* → EST003 | This study |
| EST083 | PolC-Halo | PY79 *polC-halo loxP spec* | Gift of Joseph Loparo (Harvard Medical School); strain ET096 | — |

| | | | | |
|---|---|---|---|---|
| EST085 | mYPet-DnaE | PY79 *loxP spec mYPet-dnaE* | Gift of Joseph Loparo (Harvard Medical School); strain ET121 | — |
| EST089 | Halo-DnaE | PY79 *loxP spec halo-dnaE* | Gift of Joseph Loparo (Harvard Medical School); strain ET138 | — |
| EST101 | *E. coli* prototrophic wild-type strain | MG1655 | — | — |
| EST111 | ΔPol Y1 | PY79 *yqjH::loxP spec* | EST111 | (7) |
| EST181 (BWX5235) | Halo-DnaE$^{AMGLA}$ | PY79 *loxP spec halo-dnaE*$^{AMGLA}$ *loxP kan* | Transformation: *dnaE*$^{AMGLA}$ *loxP kan* → EST089 | This study |
| EST183 (BWX5237) | Halo-DnaE$^{QLSLF}$ | PY79 *loxP spec halo-dnaE*$^{QLSLF}$ *loxP kan* | Transformation: *dnaE*$^{QLSLF}$ *loxP kan* → EST089 | This study |
| EST185 (BWX5240) | Halo-DnaE$^{AMGLA}$ DnaX-mYPet | PY79 *loxP spec halo-dnaE*$^{AMGLA}$ *loxP kan dnaX-mYpet cat Ω pWX340a* | Transformation: EST181 → EST053 | This study |
| EST187 (BWX5242) | Halo-DnaE$^{QLSLF}$ DnaX-mYPet | PY79 *loxP spec halo-dnaE*$^{QLSLF}$ *loxP kan dnaX-mYpet cat Ω pWX340a* | Transformation: EST183 → EST053 | This study |
| EST203 | Halo-DnaE DnaX-mYPet | PY79 *loxP spec halo-dnaE dnaX-mYpet cat Ω pWX340a* | Transformation: EST089 → EST053 | This study |
| EST205 | *E. coli* protein expression strain | BL21(DE3) | — | — |
| EST281 | DnaX-mYPet (ectopic) | PY79 *yvbJ::dnaX-mYpet cat* | Transformation: *yvbJ:dnaX-mYPet cat* → EST003 | This study |
| EST283 (BWX5908) | PolC$^{QMGLF}$ | PY79 *polC*$^{QMGLF}$ *loxP spec* | Transformation: *polC*$^{QMGLF}$ *loxP spec* → EST003 | This study |
| EST285 (BWX5934) | PolC$^{QMGLF}$-mYPet | PY79 *polC*$^{QMGLF}$*-mYPet cat* | Transformation: *polC*$^{QMGLF}$*-mYPet cat* → EST003 | This study |
| EST287 | PolC-Halo DnaX-mYPet (ectopic) | PY79 *polC-halo loxP spec yvbJ::dnaX-mYpet cat* | Transformation: EST083 → EST281 | This study |

**Table S14.** Imaging dataset size.

| **Dataset/Condition** | **Figure(s)** | **Number of Days** | **Number of Replicates** | **Number of Cells** | **Number of Tracks or Foci** |
|---|---|---|---|---|---|
| $DnaE^{QMGLF(WT)}$ $D^*$ | 4C, S5A | 8 | 8 | 2,391 | 21,192 |
| $DnaE^{AMGLA}$ $D^*$ | 4C | 6 | 6 | 2,076 | 16,808 |
| $DnaE^{QLSLF}$ $D^*$ | 4C | 6 | 6 | 1,849 | 11,226 |
| $PolC^{QLSLF(WT)}$ $D^*$ | 4C | 3 | 4 | 959 | 14,522 |
| $DnaE^{QMGLF(WT)}$-DnaX $g(r)$ | 4D | 8 | 8 | 2,391 | 21,753 |
| $PolC^{QLSLF(WT)}$-DnaX $g(r)$ | 4D | 3 | 4 | 959 | 14,987 |
| $DnaE^{AMGLA}$-DnaX $g(r)$ | 4D | 6 | 6 | 2,076 | 17,843 |
| $DnaE^{QLSLF}$-DnaX $g(r)$ | 4D | 6 | 6 | 1,849 | 11,909 |
| $PolC^{QLSLF(WT)}$-mYPet cellular localization | 5C | 3 | 3 | 482 | 823 |
| $PolC^{QMGLF}$-mYPet cellular localization | 5C | 3 | 3 | 671 | 1,194 |
| False positive spots $D^*$ | S5B | 3 | 3 | 724 | 84 |
| False positive spots localization | S5B | 3 | 3 | 724 | 93 |
| DnaX cellular localization<br>Endogenous fusion | S6A | 3 | 3 | 724 | 1,175 |
| DnaX cellular localization<br>Ectopic fusion | S6B | 2 | 2 | 711 | 991 |
| DnaX cellular localization<br>Ectopic fusion + $PolC^{QMGL(WT)}$-Halo | S6C | 3 | 4 | 959 | 1,357 |
| DnaX cellular localization<br>Endogenous fusion + Halo-$DnaE^{QMGLF(WT)}$ | S6D | 8 | 8 | 2,391 | 3,813 |
| DnaX cellular localization<br>Endogenous fusion + Halo-$DnaE^{AMGLA}$ | S6E | 6 | 6 | 2,076 | 3,539 |
| DnaX cellular localization<br>Endogenous fusion + Halo-$DnaE^{QLSLF}$ | S6F | 6 | 6 | 1,849 | 2,281 |
| $DnaE^{QMGLF(WT)}$ jump lengths | S7 | 8 | 8 | 2,391 | 88,030 |
| $DnaE^{AMGLA}$ jump lengths | S7 | 6 | 6 | 2,076 | 75,237 |
| $DnaE^{QLSLF}$ jump lengths | S7 | 6 | 6 | 1,849 | 53,824 |
| $PolC^{QLSLF(WT)}$ jump lengths | S7 | 3 | 4 | 959 | 61,859 |
| Halo-$DnaE^{QMGLF(WT)}$ cellular localization | S8A | 8 | 8 | 2,391 | 23,138 |
| Halo-$DnaE^{QLSLF}$ cellular localization | S8B | 6 | 6 | 1,849 | 12,474 |
| Halo-$DnaE^{AMGLA}$ cellular localization | S8C | 6 | 6 | 2,076 | 18,472 |
| $PolC^{QLSLF(WT)}$-Halo cellular localization | S8D | 3 | 4 | 959 | 15,971 |

### **Supplementary Methods**

*Overview of strain construction strategy:*

Plasmids were constructed using standard molecular biology methods. Introduction of chromosomal genetic modifications by transformation of genomic DNA or double-stranded DNA (dsDNA) fragments assembled via Gibson assembly(8) was performed exactly as described previously.(7) Tables S11 – S13 list all oligonucleotides, plasmids, and bacterial strains used in this study. Construction details for all new strains are summarized below.

Expression plasmids for *E. coli* and *B. subtilis* DnaN were cloned in the pET-28b vector. Constructs contained an N-terminal hexahistidine tag ($His_6$) and short flexible linkers (MGSSHHHHHHSSG) preceding the N-terminus of DnaN.

A weak-binding *polC* allele (*polC*$^{QMGLF}$) was generated by mutating the wild-type (WT) clamp-binding motif (CBM), QLSLF, to the *dnaE* CBM, QMGLF. Tight-binding and binding-deficient *dnaE* alleles were generated by mutation of the WT CBM, QMGLF, to the PolC CBM, QLSLF, and the double-alanine mutant, AMGLA, respectively (yielding the *dnaE*$^{QLSLF}$ and *dnaE*$^{AMGLA}$ alleles).

PolC and DnaE fusions to the self-labeling HaloTag(9) were constructed for single-molecule fluorescence microscopy. A C-terminal PolC-Halo fusion was built with an 11 amino acid linker (GSGQGSGPGSG) between the PolC C-terminus and the HaloTag N-terminus. Halo-DnaE constructs for microscopy were N-terminal fusions with an 8 amino acid linker (LEGSGQGP). Halo-DnaE fusions bearing the tight-binding and binding-deficient CBM mutations were also constructed. As a replisome marker, the clamp-loader complex subunit DnaX was fused at its C-terminus to the yellow fluorescent protein (YFP) variant mYPet(10) with an 8 amino acid linker (LEGSGQGP). Because the PolC-Halo fusion was synthetically lethal in combination with a DnaX-mYPet fusion at the endogenous *dnaX* locus, we created an ectopic DnaX-mYPet fusion introduced at the locus of the non-essential gene *yvbJ*; this gene plays no known role in replication. Replisomes in this merodiploid strain are expected to contain a mix of tagged and untagged DnaX molecules. Addition of the PolC-Halo fusion to this strain was tolerated, consistent with prior reports of strains bearing endogenous PolC fusions and ectopic DnaX fusions.(11, 12)

Although the PolC-Halo fusion was synthetically lethal with the *polC*$^{QMGLF}$ weak-binding CBM mutation, a corresponding C-terminal mYPet fusion was viable. Therefore, PolC-mYPet and PolC$^{QMGLF}$-mYPet fusions were also constructed for microscopy. These fusions contained the 8 aa linker (LEGSGQGP).

*Detailed plasmid construction information:*

**pEST007:** pET-28b-$His_6$-Ec-DnaN. The pET-28b backbone was amplified from plasmid pEST005 using oligonucleotides oEST086 and oEST087. The *E. coli dnaN* gene was amplified from strain EST101 using oligonucleotides oEST082 and oEST083. The two fragments were joined by Gibson assembly and transformed into strain EST001.

**pEST009:** pET-28b-$His_6$-Bs-DnaN. The pET-28b backbone was amplified from plasmid pEST005 using oligonucleotides oEST086 and oEST087. The *B. subtilis dnaN* gene was amplified from strain EST003 using oligonucleotides oEST084 and oEST085. The two fragments were joined by Gibson assembly and transformed into strain EST001.

*Detailed strain construction information:*

**BWX2231:** PolC WT spec. The region upstream of the *polC* gene was amplified from strain EST003 using oligonucleotides oWX916 and oWX919, the *loxP-spec-loxP* cassette was amplified from plasmid pWX466 using universal primers oWX438 and oWX439, and the region downstream of the *polC* gene was amplified from strain EST003 using oligonucleotides oWX920 and oWX921. The three fragments were joined by Gibson assembly and transformed into strain EST003.

**EST007:** SSB Δ35. The region upstream of the *ssb* gene was amplified from strain EST003 using oligonucleotides oWX2235 and oWX2238, the *loxP-kan-loxP* cassette was amplified from plasmid pWX470 using universal primers oWX438 and oWX439, and the region downstream of the *ssb* gene was amplified from EST003 using oligonucleotides oWX2239 and oWX2240. The three fragments were joined by Gibson assembly and transformed into strain EST003.

**EST009:** DnaE loxP kan. The region upstream of the *dnaE* gene was amplified from strain EST003 using oligonucleotides oWX900 and oWX903, the *loxP-kan-loxP* cassette was amplified from plasmid pWX470 using universal primers oWX438 and oWX439, and the region downstream of the *dnaE* gene was amplified from strain EST003 using oligonucleotides oWX904 and oWX905. The three fragments were joined by Gibson assembly and transformed into strain EST003.

**EST013:** DnaE-AMGLF. The upstream and N-terminal part of *dnaE* was amplified from strain EST003 using oligonucleotides oWX900 and oWX2241, and the region containing the C-terminal part of *dnaE*, the *loxP kan* cassette, and the downstream of *dnaE* was amplified from strain EST009 using oligonucleotides oWX905 and oWX2242. The three fragments were joined by Gibson assembly and transformed into strain EST003.

**EST043:** $DnaE^{AMGLA}$. The upstream and N-terminal part of *dnaE* was amplified from strain EST013 using oligonucleotides oWX900 and oWX2354, and the region containing the C-terminal part of *dnaE*, the *loxP kan* cassette, and the downstream of *dnaE* was amplified from strain EST013 using oWX905 and oWX2355. The two fragments were joined by Gibson assembly and transformed into strain EST003.

**EST047:** DnaE$^{QLSLF}$. The upstream and N-terminal part of *dnaE* was amplified from strain EST003 using oligonucleotides oWX900 and oWX2249, and the region containing the C-terminal part of *dnaE*, the *loxP kan* cassette, and the downstream of *dnaE* was amplified from strain EST009 using oligonucleotides oWX905 and oWX2357. The two fragments were joined by Gibson assembly and transformed into strain EST003. The transformants were PCR amplified using oWX902 and oML83 and sequenced using oWX902.

**EST053:** DnaX-mYPet. Plasmid pWX340 was transformed into strain EST003 and integrated in to the chromosome by single crossover.(3)

**EST057:** PolC-mYPet. The C-terminal domain of *polC* was amplified from strain EST003 using oligonucleotides oWX225 and oWX323, and the downstream region of *polC* was amplified from strain EST003 using oligonucleotides oWX324 and oWX228. These two PCR products were used as primers to amplify the *mYpet cat* allele from plasmid pWX318 according to the long-flanking-homology (LFH) method.(13) The final PCR product was transformed into strain EST003.

**EST181:** Halo-DnaE$^{AMGLA}$. The DnaE$^{AMGLA}$ construct, the *loxP kan* cassette, and the downstream region of *dnaE* were amplified from strain EST043 using oligonucleotides oWX900 and oWX905. The PCR product was transformed into strain ET089.

**EST183:** Halo-DnaE$^{QLSLF}$. The DnaE$^{QLSLF}$ construct, the *loxP kan* cassette, and the downstream region of *dnaE* were amplified from strain EST047 using oligonucleotides oWX900 and oWX905. The PCR product was transformed into strain ET089.

**EST185:** Halo-DnaE$^{AMGLA}$ DnaX-mYPet. The *loxP spec halo-dnaE$^{AMGLA}$* allele was transferred from strain EST181 to strain EST053 by transformation with genomic DNA.

**EST187:** Halo-DnaE$^{QLSLF}$ DnaX-mYPet. The *loxP spec halo-dnaE$^{QLSLF}$* allele was transferred from strain EST183 to strain EST053 by transformation with genomic DNA.

**EST203:** Halo-DnaE DnaX-mYPet. The *loxP spec halo-dnaE* allele was transferred from strain EST089 to strain EST053 by transformation with genomic DNA.

**EST281:** DnaX-mYPet (ectopic). The N-terminal 500 bp of *yvbJ* and the upstream were amplified from strain EST003 using oligonucleotides oEST118 and oEST119. The *dnaX-mYPet* fusion and the chloramphenicol cassette, including 150 bp of the *dnaX* upstream, were amplified from strain EST053 using oligonucleotides oEST120 and oEST121. The C-terminal 500 bp of *yvbJ* and the downstream were amplified from strain EST003 using oligonucleotides oEST122 and oEST123. The three fragments were joined by Gibson assembly and transformed into strain EST003.

**EST283:** PolC$^{QMGLF}$. The region upstream of the *polC* gene was amplified from strain EST003 using oligonucleotides oWX916 and oWX3633, and the linker, *loxP spec* cassette, and the region downstream of *polC* were amplified from strain BWX2231 using oligonucleotides oWX921 and oWX3634. The two fragments were joined by Gibson assembly and transformed into strain EST003.

**EST285:** PolC$^{QMGLF}$-mYPet. The region upstream of the *polC* gene was amplified from strain EST003 using oligonucleotides oWX916 and oWX3633, and the linker, mYPet, cat cassette, and region downstream of *polC* were amplified from strain EST057 using oligonucleotides oWX921 and oWX3692.

**EST287:** PolC-Halo DnaX-mYPet (ectopic). The *polC-halo loxP spec* allele was transferred from strain EST083 to strain EST281 by transformation with genomic DNA.
